# Decoding the Cure-all Effects of Ginseng

**DOI:** 10.1101/2023.04.05.535784

**Authors:** Shining Loo, Antony Kam, Bamaprasad Dutta, Xiaohong Zhang, Nan Feng, Siu Kwan Sze, Chuan-Fa Liu, Xiaoliang Wang, James P. Tam

## Abstract

Ginseng has been known as a “cure-all” traditional medicine to treat various illnesses and as an adaptogen to relieve stress. However, the known active compounds of ginseng are small-molecule metabolites. Here we report ginsentides, which are disulfide-dense, super-stable and cell-penetrating peptides with 31–33 amino acids, as active compounds and adaptogens that restore homeostasis in response to stress. Using mass spectrometry-based target identification and functional studies, we show that ginsentides promote vasorelaxation by producing nitric oxide through endothelial cells via the PI3K/Akt signaling pathway. Ginsentides were also found to alleviate α1-adrenergic receptor overactivity by reversing phenylephrine-induced constriction of the aorta, decrease monocyte adhesion to endothelial cells via CD166/ESAM/CD40, inhibit P2Y12 receptors, reduce platelet aggregation, and thrombus formation in the lung. Orally administered ginsentides were effective in anti-stress behavior using animal models of tail suspension and forced swimming tests. Together, these results suggest that ginsentides interact with multiple systems to restore homeostasis by reversing stress-induced physiological changes and provide new insights into the panacea medicinal effects of ginseng.

## Introduction

Ginseng is a popular traditional medicine with a wide range of pharmacological effects and a long history of usage for health maintenance and treatment of diseases^1^. Numerous clinical studies using ginseng extracts have shown that ginseng improves immune functions and cognitive and physical performance and can be used to treat cancer and cardiovascular diseases^1–4^.

Ginseng is the collective name for thirteen species in the Panax genus in the Araliaceae family. Among them, *Panax ginseng* (Asian ginseng), *Panax quinquefolius* (American ginseng), and *Panax notoginseng* (sanqi) have been extensively studied for their panacea or “cure-all” medicinal effects^5^. To date, more than 200 chemicals have been identified from *P. ginseng*^6–8^. These include various small-molecule metabolites, such as triterpenes, saponins, polyacetylenes, alkaloids, phenolics, and biopolymers such as polysaccharides, peptidoglycans, and proteins^6–8^. Currently, ginsenosides, a class of steroidal glycosides and triterpene saponins, are thought to be the principal active compounds in ginseng^1, 3, 9^. Specifically, two major ginsenosides, the protopanaxdiol-type Rb1 and the protopanaxtriol-type Rg1, have been reported to elicit anti-inflammatory and cardioprotective activities, partially through the androgen, estrogen, and glucocorticoid receptors^10–13^. However, other families of natural products conferring the cure-all or panacea effects of *P. ginseng* remain to be discovered.

Peptides are conspicuously absent among ginseng’s families of bioactive compounds. A common belief in natural product drug discovery is that peptides are too labile to heat and proteases to have significant therapeutic value, especially when taken orally. However, heavily cross-linked peptides, known as cysteine-rich peptides (CRPs), could be exceptions due to their high tolerance to heat and proteolytic degradation^14–17^. Recently, we discovered ginsentides, which are glycine- and cysteine-rich peptides with 31–33 amino acids and four disulfide bonds^18^. The disulfide arrangement in ginsentides is different from all known plant CRPs and forms a highly constrained pseudocyclic, with cysteine flanking both the N- and C-termini that form loops with the internal cysteine^18^. As a result, pseudocyclic ginsentides are highly resistant to proteolytic degradation^18^. Representative examples include ginsentide TP1, TP3, and TP8, from *Panax ginseng* and *Panax notoginseng*, *Panax ginseng*, and *Panax quinquefolius*, respectively, are all 31 amino acid residues long, highly homologous and found abundantly in ginseng plants (**Fig.1**).

**Fig. 1.**
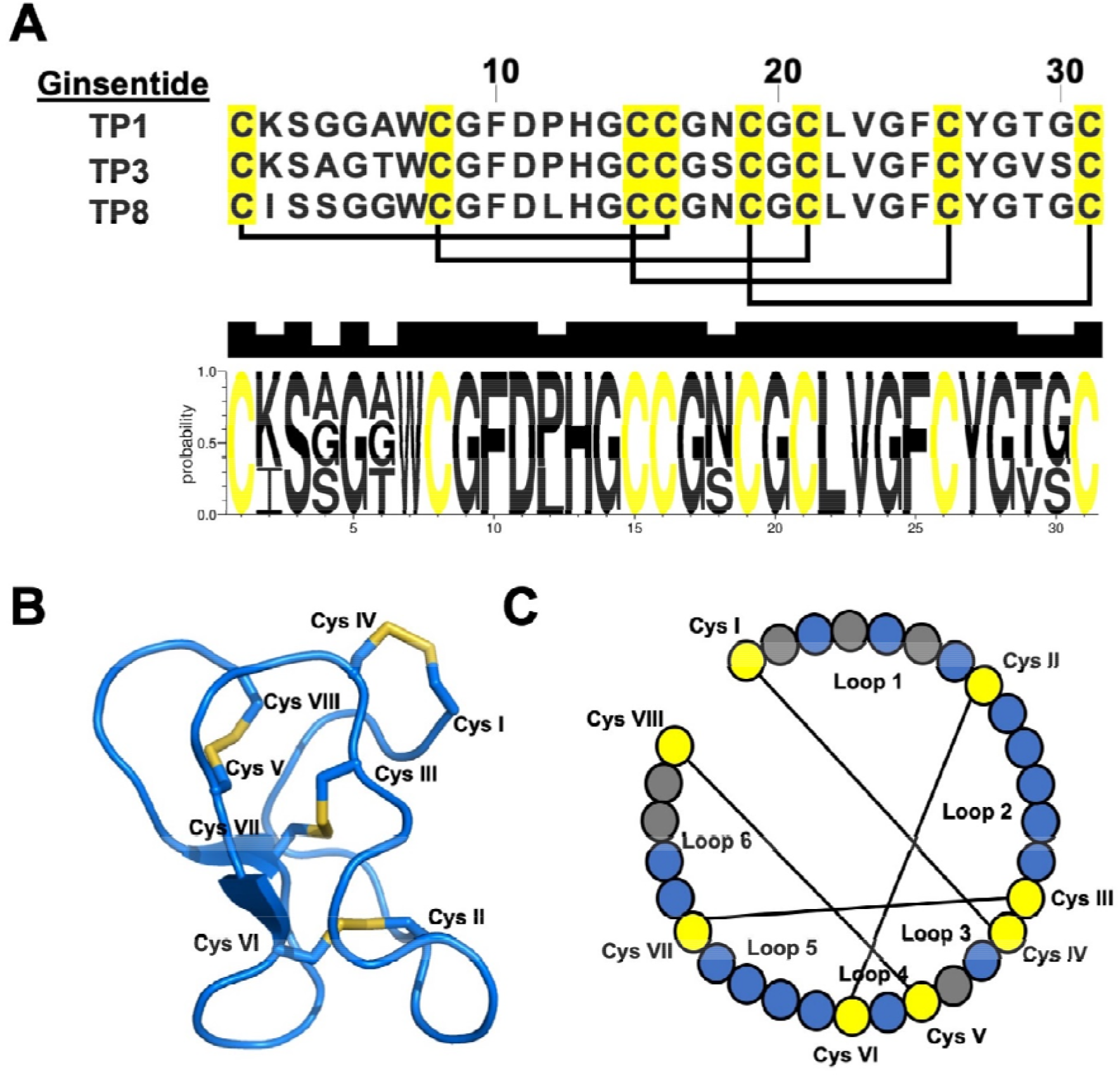
(A) Primary amino acid sequences of ginsentide TP1 (*Panax ginseng* and *P. notoginseng*), TP3 (*P. ginseng*), and TP8 (*P. quinquefolius*). Web logo showing the highly conserved and homologous amino acid residues of ginsentides TP1, TP3, and TP8. (B) The tertiary structure of ginsentide TP1 (PDB: 2ML7). (C) Illustration of the primary amino acid sequence of TP1, TP3, and TP8, with the colored circles representing amino acid residues (yellow: cysteine; blue: conserved amino acid residues; gray: unconserved amino acid residues) and lines representing disulfide linkages.

The panacea effect of ginseng, as described in ancient texts such as “The Divine Farmer’s Materia Medica” and “The Yellow Emperor’s Classic of Internal Medicine”, includes medicinal benefits like invigorating and restoring cellular homeostasis by revitalizing visceral organs, stimulating rapid recovery from illnesses, improving blood circulation, counteracting the effects of physical and emotional stress, relieving fatigue, and improving stamina^19–22^. Consequently, ginseng is widely prescribed to treat many diseases. An example is the highly influential “Treatise on Febrile Diseases”, which documents 113 classic formulations, of which at least 20 contain ginseng^23^. A modern-day interpretation of ginseng’s medicinal value is that it is a general tonic for overall health, boosts the immune system, helps prevent illness, and improves overall physical and mental stress adaptation^24–27^. Because of its ability to modulate the body’s stress response and restore balance, ginseng is also an adaptogen^28^. However, the current known active compounds based on ginsenosides cannot adequately explain the panacea effects of ginseng.

Here, we show that ginsentides, a novel family of bioactive peptides, are adaptogns by eliciting diverse physiological functions that confer cardiovascular benefits, contributing to the anti-stress and panacea effects attributed to ginseng in traditional medicine.

## Results

### Preparation of ginsentides TP1, TP3, and TP8 from three common ginseng plants

For target identification, validation, and functional studies, we selected the three most abundant ginsentides: TP1, found in *P. ginseng* and *P. notoginseng*, TP3, found in *P. ginseng* and *P. quinquefolius*, and TP8 from *P. quinquefolius*. All three have highly conserved sequences, with TP1 differing from TP3 and TP8 by five and two amino acid residues, respectively. Previously, we showed that the distributions of ginsentides are species and tissue-dependent^18^. We performed large-scale extraction and purification of ginsentide TP1 from *P. notoginseng* flowers, TP8 from *P. quinquefolius* flowers, and TP3 from *P. ginseng* seeds. The crude aqueous extracts were fractionated using C18 reversed-phase and strong cation-exchange flash chromatography, and the presence of ginsentides was monitored by mass spectrometry. The concentrated fractions were further purified by reversed-phase high-performance liquid chromatography (RP-HPLC) to achieve a single peak with over 95% purity (**Supplementary data S1**). From 1 kg of ginseng flowers or seeds, we obtained approximately 200 mg of ginsentides TP1, TP3, or TP8.

In addition to isolating ginsentides from plant materials, we successfully obtained ginsentide TP1 via total chemical synthesis. A synthetic ginsentide precursor was prepared using a stepwise solid-phase synthesis method. After TFA cleavage to remove the protecting groups and release it from the resin support, the S-reduced linear precursor was immediately subjected to oxidative folding under anaerobic conditions, followed by purification using RP-HPLC. The purified synthetic ginsentides, such as ginsentide TP1, were identical to their native form as determined by RP-HPLC, MS, and 1D and 2D NMR. This suggests that these synthetic ginsentides are chemically identical to native ginsentides and that their disulfides are correctly folded (**Supplementary data S2**).

### Ginsentide TP1 is a cell-penetrating peptide

Our previous studies showed that certain structural-compact and hydrophobic plant-derived cysteine-rich peptides can internalize into cells to interact with intracellular protein targets^17, 29^. To investigate whether the hydrophobic and structural-constrained ginsentides are cell-penetrating, we prepared fluorescent-labeled Cy3-TP1 using cyanine 3 (Cy3)-N-hydroxysuccinimide ester at its lysine sidechain. **Fig. 2A** shows that Cy3-TP1 was internalized into HUVEC-CS cells using confocal microscopy. The cellular uptake of Cy3-TP1 at 4 °C was substantially reduced compared to 37 °C using flow cytometry (**Fig. 2D**). Pre-treating cells with endocytosis inhibitors reduced Cy3-TP1 uptake (**Fig. 2E**). These results suggest that the cellular uptake of Cy3-TP1 involves energy-dependent endocytosis and ginsentide TP1 may have intracellular targets.

**Fig. 2.**
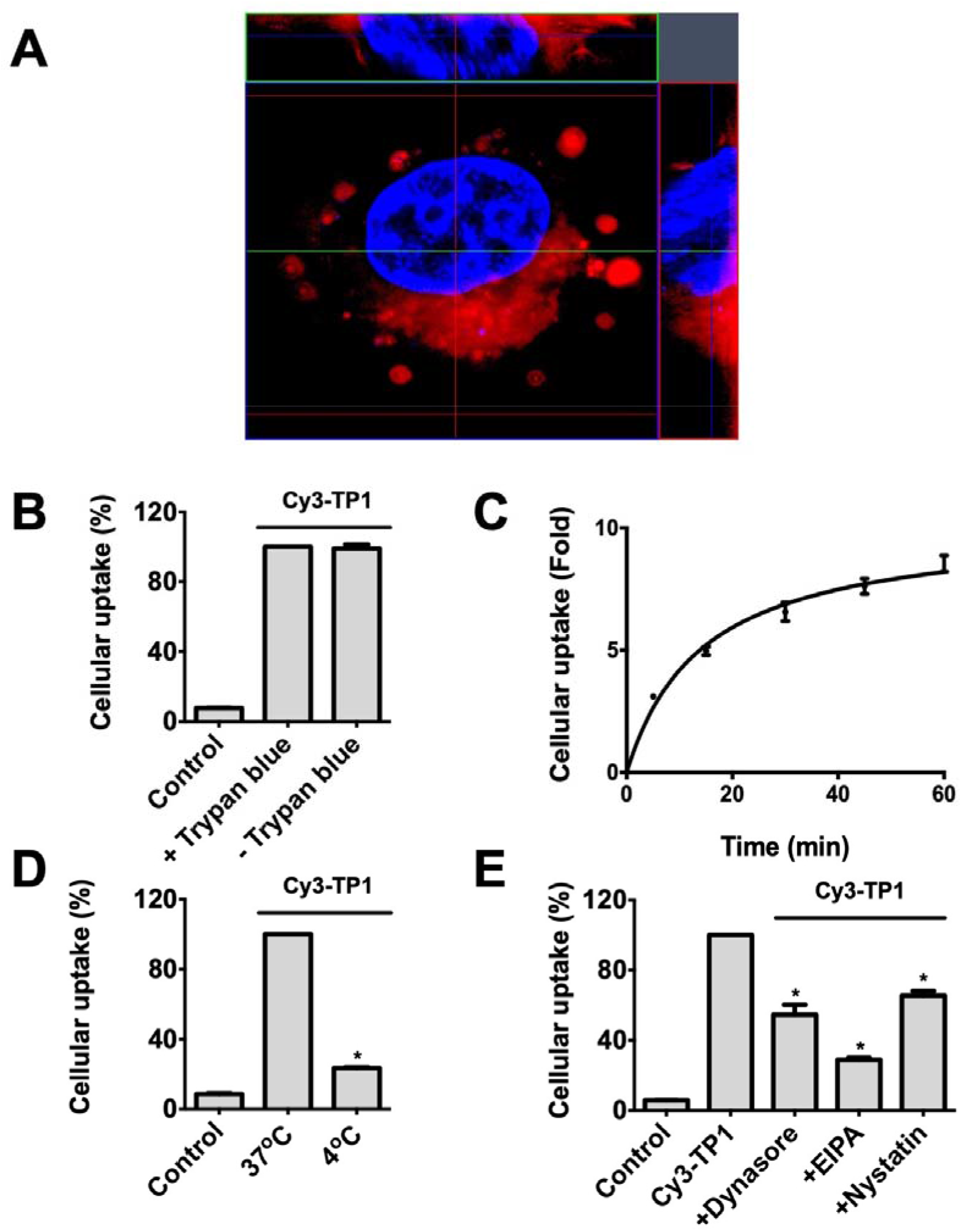
Cy3-TP1 is cell-penetrating. (A) Z-stack image of Cy3-TP1 in HUVEC-CS cells using confocal microscopy. Using flow cytometry, the cellular uptake of Cy3-TP1 into HUVEC-CS cells was (B) unaffected by trypan blue, (C) time-dependent, (D) temperature dependent, and (E) affected by endocytosis inhibitors (Dynasore, EIPA, and Nystatin). All results are expressed as mean ± S.E.M. from three separate experiments. *p < 0.05 compared to control.

### Proteomic profiling and target identification of intra- and extracellular proteins of ginsentide TP1

To identify the intra- and extra-cellular targets of ginsentide TP1, we used affinity-enrichment MS profiling. Again, we immobilized ginsentide TP1 at the lysine sidechain with UltraLink Neutravidin agarose resins for target identification. Three separate experiments were performed, and the samples were tested in triplicate. A pull-down assay followed by LC-MS/MS analysis revealed that 256 proteins were uniquely identified in the ginsentide TP1 pull-down samples (**Figure 3**) (**Dataset 1** and **2**), after removing the background proteins found in parallel control experiments. Bioinformatics analysis of the ginsentide TP1 interactome data suggested the involvement of several important biological pathways, including PIP3 activation, Akt signaling, platelet homeostasis, the MAPK family signaling cascade, interleukin-1 signaling, mTORC1-mediate signaling, L1CAM interactions, and cellular stress responses (**Dataset 3**).

**Fig. 3.**
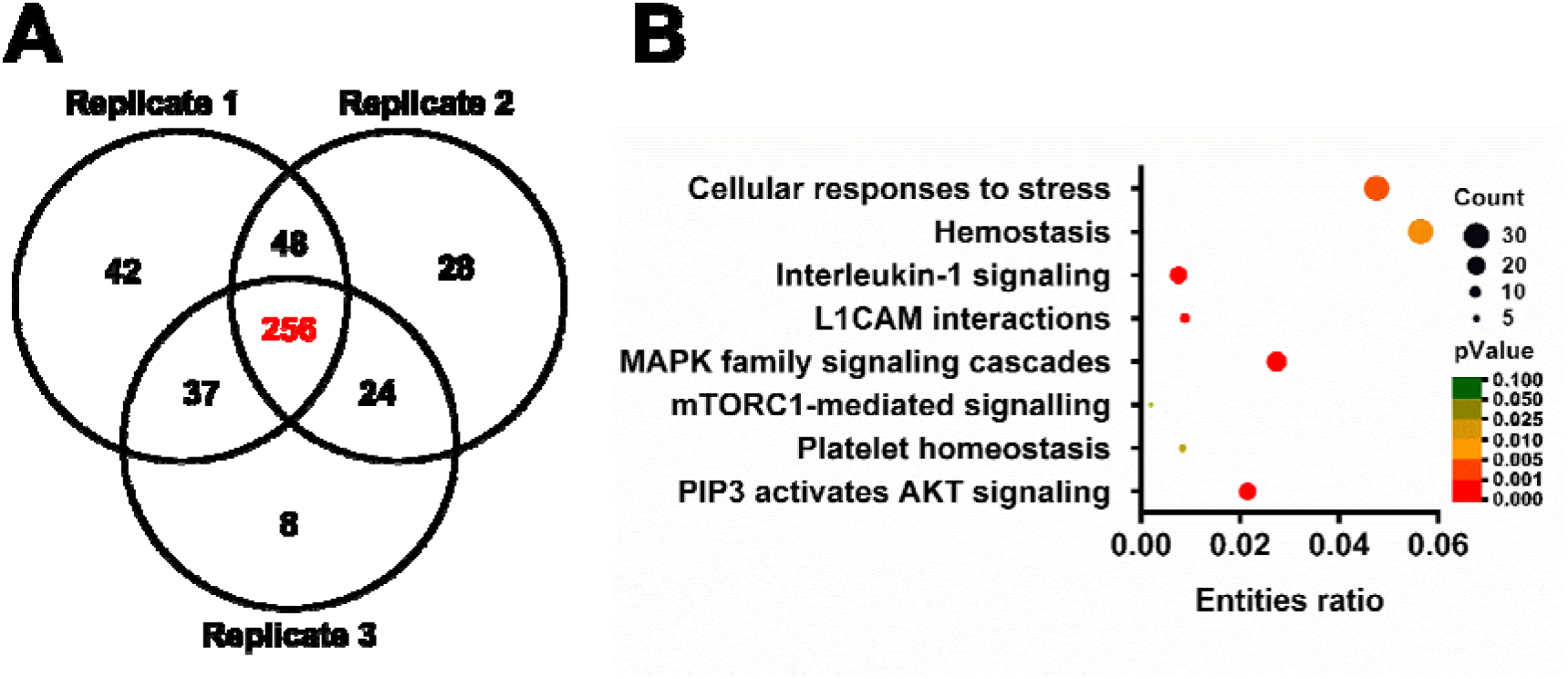
Proteomic profiling of intra- and extra-cellular targets of Ginsentide TP1. (A) Venn diagram of the uniquely identified proteins for ginsentide TP1 from three experimental replicates. (B) Enriched interacting pathways of ginsentide TP1. Online bioinformatics tool Reactome (https://reactome.org/) was used for the pathway analysis.

### Ginsentides produce endothelium-derived nitric oxide through PI3K/Akt/eNOS

The cardiovascular effects of ginseng extracts have been extensively studied, and several reports have shown that they have vasodilatory and blood pressure-lowering effects by increasing nitric oxide (NO) levels^30–32^. NO is a crucial signaling molecule that expands the blood vessels and maintains proper vascular tone. The Akt pathway controls the regulation of NO production^33^. Supported by our proteomic analyses that ginsentide TP1 affects Akt signaling, we examined the effects of ginsentide TP1 on NO production in endothelial cells and its relationship with the Akt signaling pathway. Our results show that ginsentides TP1 (*P. ginseng* and *P. notoginseng*), TP3 (*P. ginseng*), and TP8 (*P. quinquefolius*) increase NO production in endothelial cells (**Fig. 4A** and **B**). Ginsentide TP1 was three times more effective than ginsenosides Rb1 and Rg1 on a molar basis (**Fig. 4A**). To eliminate the possibility of ginsenoside contamination in our naturally derived ginsentides, we showed that the synthetic ginsentide TP1 exerts similar effects as its native form (**Fig. 4C**).

**Fig. 4.**
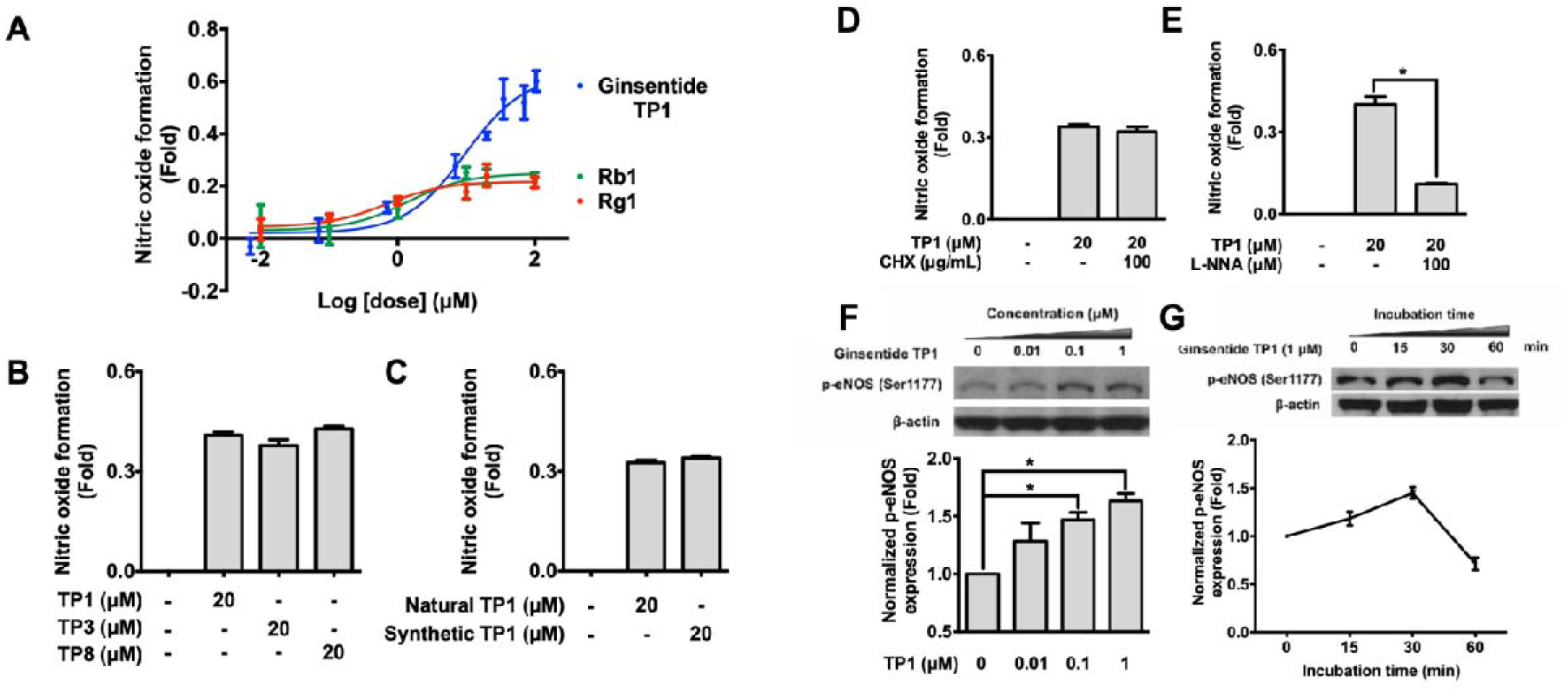
Ginsentides promote nitric oxide (NO) formation from endothelial cells. (A) HUVEC-CS cells were exposed to ginsentide TP1 and ginsenosides Rg1 and Rb1 for 1 h. (B) HUVEC-CS cells were exposed to ginsentides TP1, TP3, and TP8 for 1 h. (C) HUVEC-CS cells were exposed to natural and synthetic ginsentide TP1 for 1 h. (D) HUVEC-CS cells were exposed to ginsentide TP1 with and without cycloheximide treatment, a protein translation inhibitor, or (E) L-N^G^-Nitroarginine (L-NNA), a competitive inhibitor of nitric oxide (NO) synthase for 1 h. Intracellular NO formation was measured with DAF-2 DA staining and expressed as normalized fluorescence intensity. Representative western blot analysis on the (F) dose-dependent effects and (G) time-dependent effects of ginsentide TP1 on phosphorylated eNOS (p-eNOS) accumulations in HUVEC-CS cells. All results are expressed as mean ± S.E.M. in three separate experiments. *p < 0.05 compared to the control.

Activating intracellular eNOS results in NO formation^34^. To confirm that ginsentide TP1-induced NO production depends on NOS activity, we pre-treated endothelial cells with L-N^G^-nitroarginine (L-NNA), a competitive inhibitor of NOS. Pre-treatment with L-NNA inhibited ginsentide TP1-induced NO production. In support, Western blot analysis showed that ginsentide TP1 induced phosphorylated eNOS (p-eNOS) accumulation in a concentration and time-dependent manner (**Figs. 4E–G**). The PI3K-Akt pathway can activate eNOS for NO formation^35^. To demonstrate the involvement of ginsentide TP1 in the PI3K-Akt pathway, we pre-treated endothelial cells with an Akt inhibitor, MK 2206, or a PI3K inhibitor, LY294002. Our results showed that both inhibitors decreased ginsentide TP1-induced NO production (**Supplementary data S3A** and **B**). Western blot analysis also showed that ginsentide TP1 induced Akt phosphorylation in a time-dependent manner (**Supplementary data S3C**). These results support that ginsentides elicit endothelial NO production through the PI3K-Akt pathway.

### Ginsentides reduce phenylephrine-induced vasoconstriction

Stress affects catecholamine release, leading to sympathetic overactivity^36, 37^. Phenylephrine, a catecholamine similar to adrenaline and a selective α1-adrenergic receptor agonist, was used to induce sympathetic nervous system activation, resulting in blood vessel constriction and increased arterial blood flow. To demonstrate that ginsentides restore the impaired homeostasis associated with sympathetic overactivity, we used isolated rat aortic rings in an ex vivo model to determine their vascular responses^38^. The aortic ring was pre-contracted with either phenylephrine or KCl. Isometric tension measurements showed that ginsentides TP1, TP3, and TP8, and synthetic TP1 reduced phenylephrine-induced but not KCl-induced aortic ring contractions (**Fig. 5**). The EC_50_ of TP1, TP3, and TP8 was approximately 0.1 µM (**Fig. 5A–C**). In contrast, ginsenosides Rb1 and Rg1 did not induce vasorelaxation when tested at concentrations up to 1 μM (**Fig. 5D and E**). Ginsentides TP1, TP3, and TP8 were selective for the α1-adrenergic receptor because they did not elicit relaxation of acetylcholine-induced bladder smooth muscle contraction and did not affect sodium and potassium ion channel activity when tested at concentrations of up to 5 µM (**Supplementary data S4-S8**). These data support that ginsentides alleviate the vascular function of α1-adrenergic receptor overactivity.

**Fig. 5.**
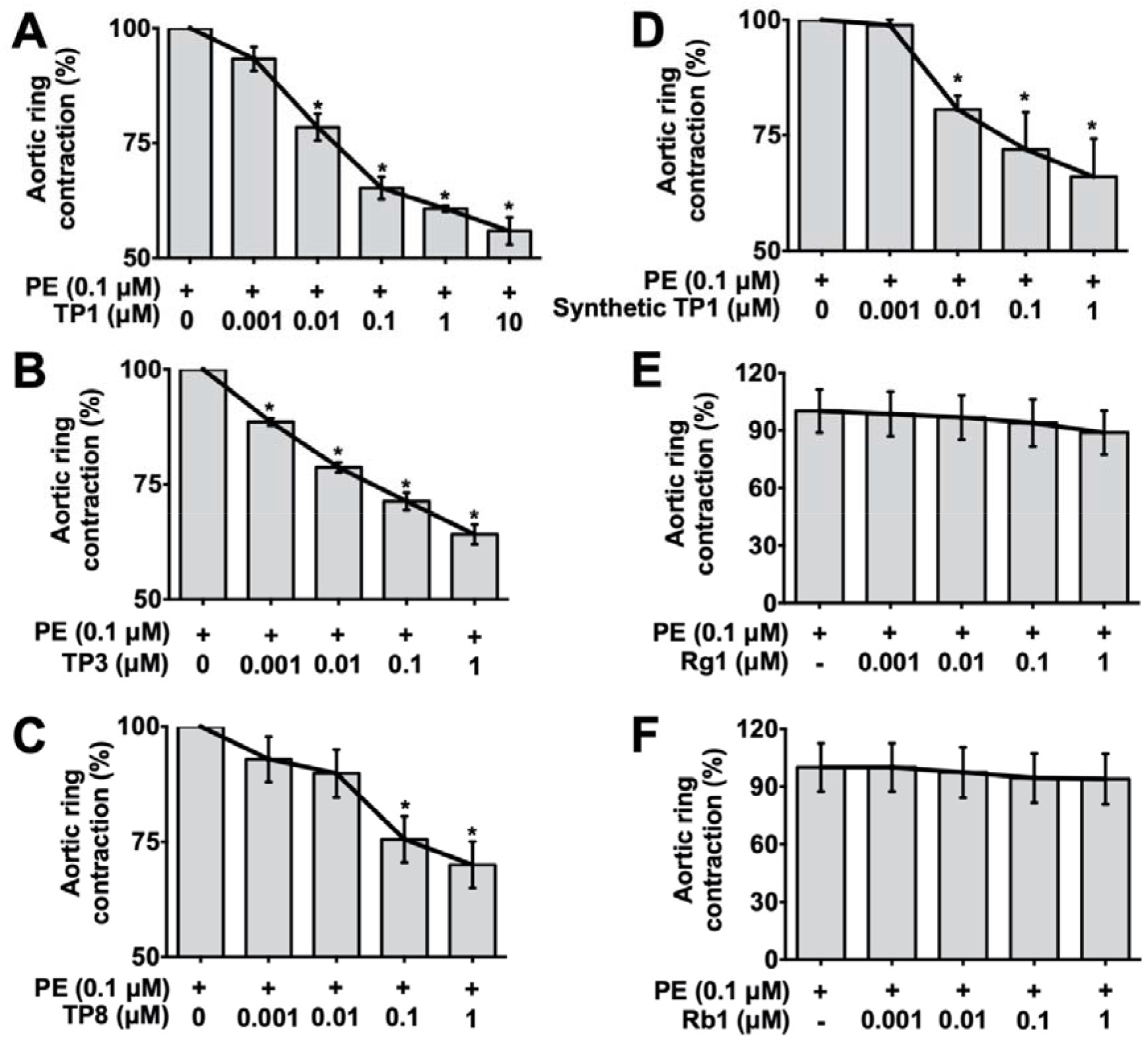
Ginsentides reduced phenylephrine-induced rat aortic ring contractions. The vascular responses of ginsentides were studied using phenylephrine (0.1 μM) pre-contracted aortic rings in isolated organ chambers. **(A)** TP1 (*P. ginseng* and *P. notoginseng*), **(B)** synthetic TP1, **(C)** TP3 (*P. ginseng*), **(D)** TP8 (*P. quinquefolius*), **(E)** ginsenoside Rg1, and **(F)** ginsenoside Rb1. The responses (isometric tension) were measured with a force-displacement transducer. Aortic ring contractions were measured in volts and are expressed as a percentage compared to controls. All results are expressed as mean ± S.E.M. from three separate experiments. *p < 0.05 compared to the control.

### Ginsentides reduce monocyte adhesion to endothelial cells

Extended periods of stress can lead to chronic inflammation, contributing to the development of cardiovascular diseases such as atherosclerosis^39–41^. One of the earliest cellular events in the development of these lesions is the adherence of cells to the endothelium^40, 41^. *P. notoginseng* is commonly used to treat cardiovascular diseases related to atherogenesis^42^. Studies have shown that ginsenosides Rg1 and Rb1 have potential anti-atherogenic effects^43, 44^. Using affinity-enrichment MS, we identified CD166 (ALCAM), CD40, and ESAM as putative targets involved in mediating cell adhesion. To support these findings, we performed pull-down assays and confirmed that ginsentide TP1 interacts with CD166 (ALCAM), CD40, and ESAM (**Fig. 6A-C**).

**Fig. 6.**
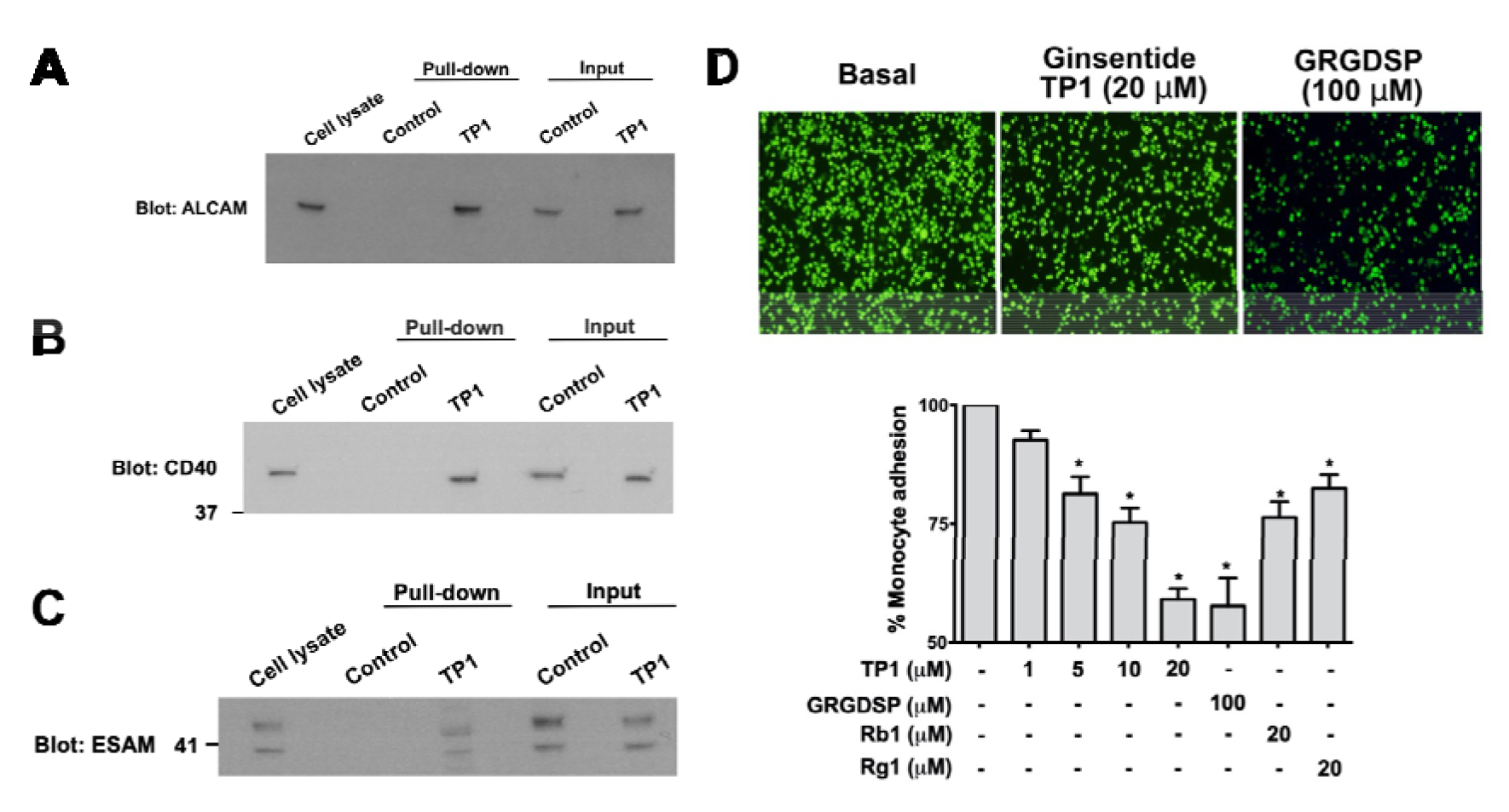
Ginsentide TP1 reduce monocyte adhesion by interacting with adhesion molecules. **(A)** Ginsentide TP1 interacts with CD166 as revealed by the pull-down western blot analyses using anti-CD166 mAb. **(B)** Ginsentide TP1 interacts with CD40 as the pull-down western blot analyses using anti-CD40 mAb. **(C)** Ginsentide TP1 interacts with ESAM as revealed by the pull-down western blot analyses using anti-ESAM mAb. **(D)** Ginsentide TP1 reduces monocyte adhesion to endothelial cells. Ginsentides TP1 and ginsenosides Rg1 and Rb1 were pre-incubated with HUVEC-CS cells for 30 min and then co-incubated with CFSE-labeled THP-1 monocytic cells for 1 h. Monocyte adhesion was measured by the fluorescence intensity of CFSE. All results are expressed as mean ± S.E.M. from three separate experiments. *p < 0.05 compared to the control.

To show that ginsentide TP1 can interfere with cell adhesion, we co-incubated ginsentide TP1 with fluorescent-labeled THP-1 monocytic cells and loaded them onto an endothelial monolayer. Ginsentide TP1 dose-dependently inhibited the basal adhesion of fluorescent-labeled THP-1 monocytic cells to the endothelial monolayer. This was compared with ginsenosides Rg1 and Rb1, which showed weaker inhibitory effects (**Fig. 6D**). The inhibitory effects of ginsentide TP1 were five times more potent than the known adhesion inhibitory peptides (GRGDSP, positive control).

### Ginsentides reduce platelet aggregation and thrombus formation

Chronic stress can activate the sympathetic nervous system, leading to the continual release of stress hormones, which may result in excessive blood clotting^36, 39^. Conditions such as atherosclerosis, deep vein thrombosis, and pulmonary embolism may occur^36, 39^. Ginseng reduces thrombotic events and improves cardiovascular health through antiplatelet mechanisms^4, 45–47^. Results from affinity-enrichment MS suggest that ginsentide TP1 affects platelet homeostasis. Additionally, ginsentide TP1 was an antagonist for the P2Y12 receptor when tested against a panel of 168 GPCR targets. **Fig. 7A** shows the dose-response curve of ginsentide TP1 inhibiting P2Y12, with an IC50 of approximately 20 µM. Conversely, ginsenosides Rb1 and Rg1 did not display inhibitory effects against P2Y12 activation at similar concentrations (**Figs. 7B and C**).

**Fig. 7.**
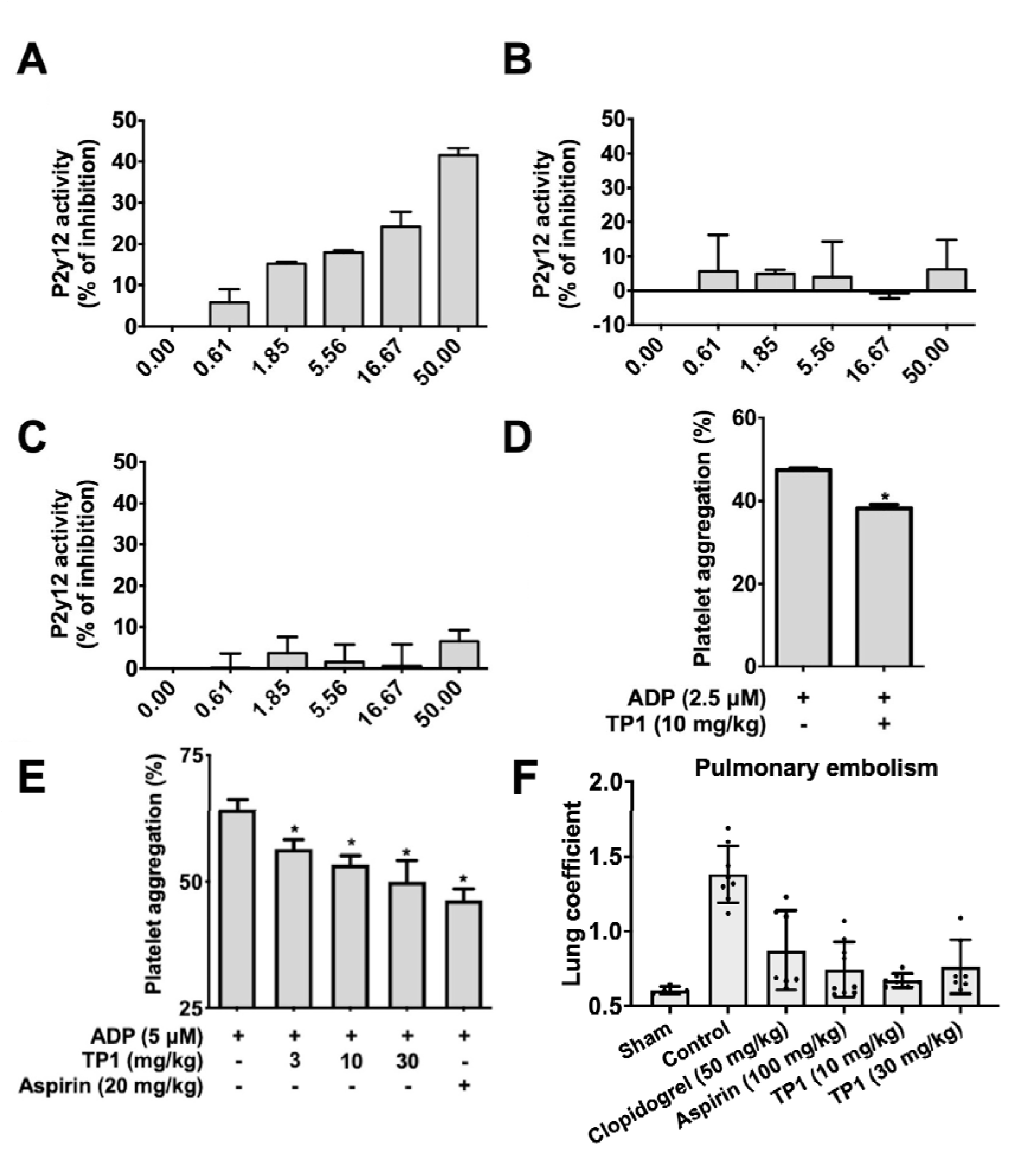
Ginsentides reduce platelet aggregation *ex vivo*. Ginsentide TP1, but not ginsenosides Rg1 and Rb1, inhibits P2Y12 activation. Effects of **(A)** ginsentide TP1, **(B)** ginsenoside Rg1, and **(C)** ginsenoside Rb1 on P2Y12 activation were analyzed with the Pathhunter eXpress β-arrestin kit (N = 2). Ginsentide TP1 (10 mg/kg) or PBS was **(D)** Intravenous injection of ginsentide TP1 (10 mg/kg) or PBS or **(E)** oral gavage of ginsentide TP1 (3, 10, 30 mg/kg) or PBS to SD rats were conducted once daily for 3 days. 30 min after the final pre-treatment, blood samples were collected, and platelets were concentrated by centrifugation. Platelet aggregation was initiated by ADP. The percentage of platelet aggregation was determined by optical density (N = 5; *p < 0.05). **(F)** Effects of oral-administered ginsentide TP1 on collagen plus adrenaline-induced acute lung thrombosis in mice.

Using an *ex vivo* platelet aggregation model, we determined the effects of ginsentide TP1 after intravenous or oral administration. Our results showed that ginsentide TP1 reduced platelet aggregation in response to ADP stimulation (**Fig. 7D** and **E**). These results also indicate that ginsentide TP1 is orally active.

We tested the anti-thrombotic effects of ginsentide TP1 *in vivo* in an acute thrombotic mouse model^48^. Acute pulmonary thrombosis was induced by tail vein injection using a combination of collagen and adrenaline^48^. Oral pre-treatment with ginsentide TP1 (10 mg/kg and 30 mg/kg) for three consecutive days significantly reduced thrombus formation in the lung compared to the saline control group. These effects were comparable to oral administration with the positive controls, clopidogrel (50 mg/kg) or aspirin (100 mg/kg) (**Fig. 7F**).

### Behavioral Tests: Tail suspension and forced swimming

Based on our experimental data, ginsentides may modulate the “fight-or-flight” response, a physiological stress response. Clinically, chronic stress can lead to depression-like behavioral changes^49, 50^. Using two well-established behavioral assays, forced swimming and tail suspension^51, 52^, we assessed the effects of ginsentide TP1 on responses to depressive-like behavior in mice. Both experiments are based on measuring the duration of immobility when the mice are exposed to an inescapable situation^51, 52^. To determine the potential antidepressant activity of ginsentide TP1, we assessed the immobility time after oral administration to mice. The forced swimming test showed that ginsentide TP1 (30 mg/kg) and the fluoxetine (20 mg/kg), positive control significantly decreased the immobility time by 71.68 ± 9.963 s and 42.52 ± 9.963 s, respectively (**Fig. 8**). Similarly, in the tail suspension test, ginsentide TP1 (30 mg/kg) and fluoxetine (20 mg/kg) as a positive control significantly decreased the immobility time by 41.38 ± 7.054 s and 30.01 ± 7.219 s, respectively (**Fig. 9**). Together, these studies suggest that ginsentide TP1 has anti-stress effects.

**Fig. 8.**
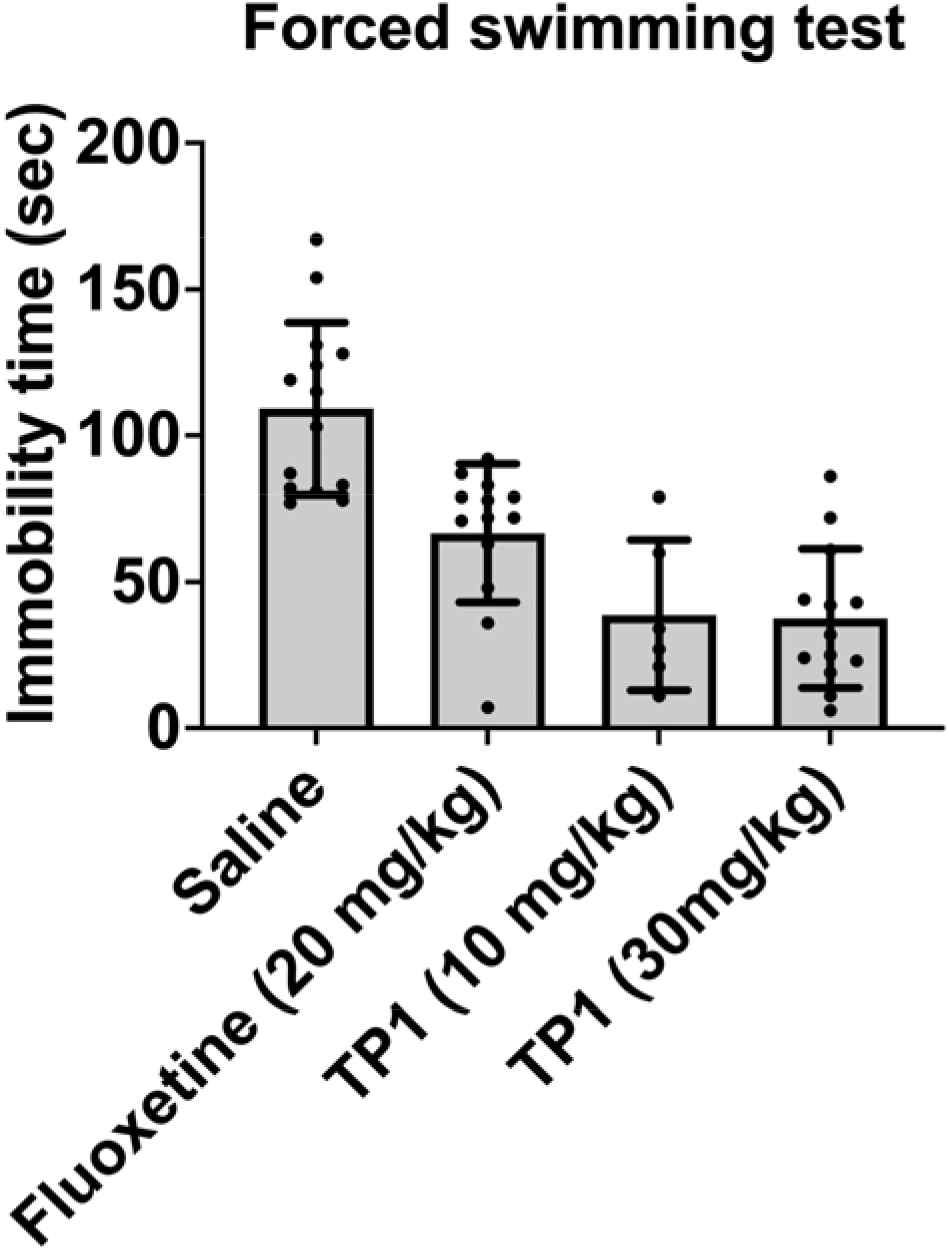
Behavioral performance of mice in the forced swimming test. Immobility period of mice in the swimming endurance stress assay measured after treatment with fluoxetine (20 mg/kg) (positive control) or different concentrations (10 or 30 mg/kg) of TP1.

**Fig. 9.**
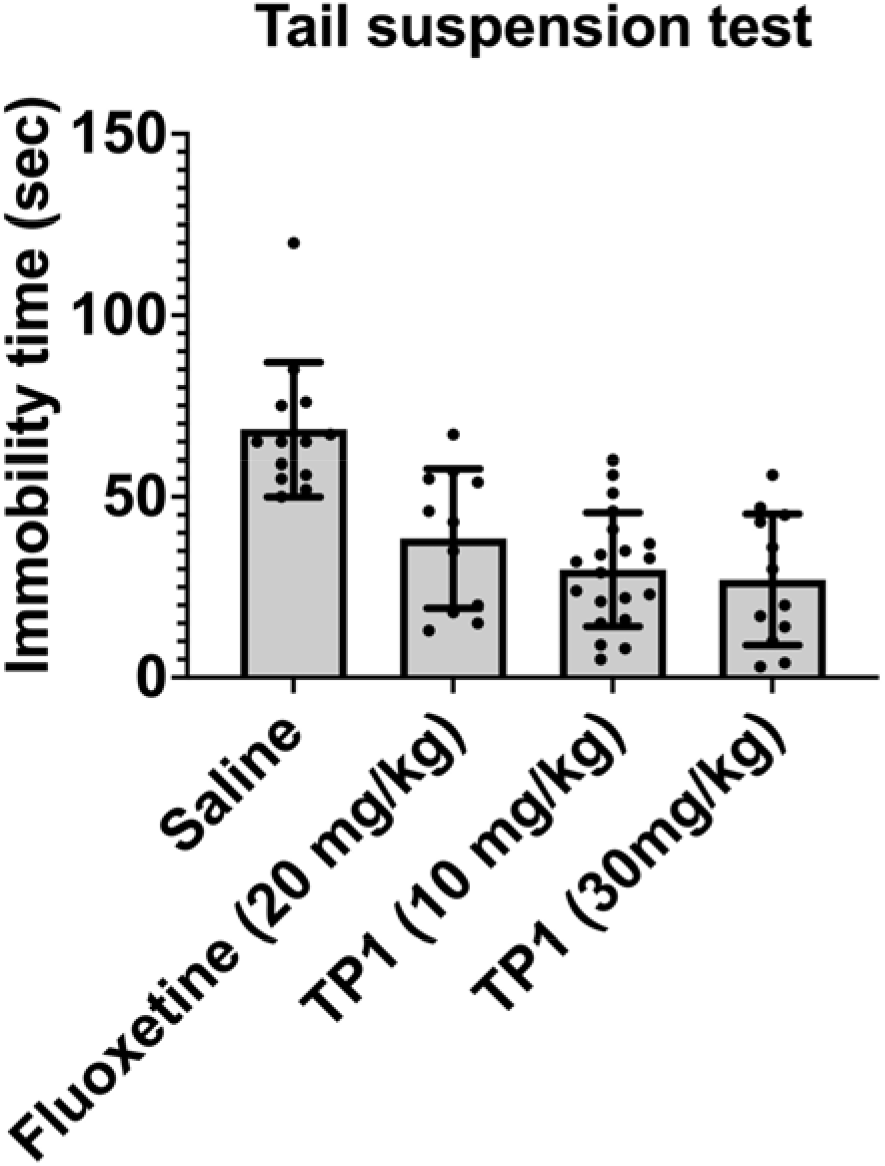
Behavioral performance of mice in the tail suspension test. Immobility period of mice in the tail suspension stress assay after oral treatment with fluoxetine (20 mg/kg) (positive control) or different concentrations (10 or 30 mg/kg) of TP1.

## Discussion

Stress is a multifaceted response to a perceived threat involving physiological and psychological changes^54^. Physiologically, the body manages stress through the “fight-or-flight” response, activating the sympathetic nervous system and releasing stress hormones like adrenaline and cortisol^55, 56^. These hormones then stimulate natural physiological reactions, including an increase in heart rate and blood pressure and activation of the coagulation cascade^56^. However, in situations such as chronic stress, erratic counteracting mechanisms may arise, resulting in physical and psychological disorders like cardiovascular diseases and depressive-like behavior^36, 39, 49, 54^. In this report, we used a combination of methods, *in vitro-*, *ex vivo-*, and *in vivo-*studies to demonstrate that ginsentides TP1, TP3, and TP8 are novel bioactive peptides and could represent major contributors to the adaptogenic effects of ginseng in relieving stress-related disorders.

To counter stress, the body uses various physiological systems, including the activation of various cardiovascular functions via multiple mechanisms^56^. *P. ginseng*, *P. quinquefolius*, and *P. notoginseng* are frequently prescribed in traditional medicine to promote blood circulation, earning them a high reputation for their cardiovascular benefits^4, 24, 44^. Specifically, we show that ginsentides increase NO production in endothelial cells, promote vasorelaxation, reduce cell adhesion and platelet aggregation, and improve depressive-like behaviors. Currently, ginsenosides are considered the major active compounds in ginseng. However, they do not fully explain the adaptogenic effects elicited by a whole-ginseng mixture.

Ginseng is generally regarded as safe for food supplements and traditional medicinal uses^25, 53^. Guided by traditional medicines, we have conducted a decade-long research program to identify orally-active super cystine-dense peptides or microproteins from medicinal plants. These microproteins, with 6-10 cysteine residues and a molecular weight of 2-4 kDa, are bioactive and show potential as therapeutic leads for cardiovascular diseases, stress, frailty, and other chronic and age-related diseases. Previously, we identified ginsentides as a group of plant-derived peptides or microproteins found in different ginseng species^18^. Currently, 14 different ginsentides have been identified. They are cystine-dense (>24% cysteine), highly homologous, 31–33 amino acids in length, with eight cysteine residues forming a unique disulfide connection^18^. Consistently, we previously showed that TP1 is non-toxic in Huh7 cells or red blood cells at concentrations up to 100 µM^18^, a concentration at least 5-fold higher than the dosages used in this paper. Furthermore, we showed that TP1 is non-immunogenic to THP-1 cells, with no observable increase in IL-6, IL-8, IL-10, and TNFα^18^.

Ginsentides TP1, TP3, and TP8 from *P. ginseng* and *P. notoginseng*, *P. ginseng*, and *P. quinquefolius*, respectively, share ∼85% homology in their primary sequences in which the eight cysteine residues, and cystiene motif are absolutely conserved. They are proteinaceous natural products. Interestingly, we found that ginsentides and ginsenosides which are small-molecule metabolites share similarity in certain bioactive functions but differ in magnitude. For example, ginsentide TP1 is more effective than ginsenosides in promoting NO production, reversing the vasoconstriction caused by phenylephrine, and reducing cell adhesion. Using GPCR profiling, we found that ginsentide TP1 can inhibit ADP-induced P2Y12 receptor activation, which could lead to reduced platelet aggregation *ex vivo* and thrombus formation *in vivo*. However, ginsenosides Rb1 and Rg1 show no activity against ADP-induced P2Y12 activation. Thus, ginsentides TP1, TP3, and TP8, represent not only the “first-in-class” active compounds in ginseng, but also a missing link in the “cure-all” use of ginseng in traditional medicines.

The identification of ginsentides as bioactive and proteolytic-stable peptides underscore the emergence of constrained peptides as active compounds in herbal medicines. Ginsentides, similar to other cysteine-rich peptides, including roseltides, bleogens, chenotides, and avenatides^14, 15, 57, 58^, are structurally constrained by multiple disulfides to render them resistant to proteolytic degradation, making them potentially orally active^14–17, 29^. Because ginsentides have a cystine-dense core, they have an “inside-out” conformation, causing the side chain of hydrophobic amino acids to be exposed^17, 29^. These exposed hydrophobic patches found in ginsentides as well as many CRPs such as roseltides from *Hibiscus sabdariffa*^17, 29^, can drive cell-penetration, allowing their internalization into cells to target intracellular targets^16, 17, 29, 57, 58^. The ability of ginsentides to target both extracellular and intracellular protein targets represent a distinctive new feature to this family of natural products.

## Conclusions

In conclusion, we demonstrated that ginsentides are orally bioactive and play a wide array of functions that provide the adaptogenic and panacea effects of ginseng, particularly for cardiovascular health. Our work greatly expands the known bioactive phytochemicals of ginseng from small molecule metabolites to non-metabolites, peptides, and microproteins and opens new avenues for ginseng research.

## Methods & Protocol

### Materials

All chemicals and solvents, unless otherwise stated, were purchased from Sigma– Aldrich (St. Louis, MO, United States) and Fisher Scientific (Waltham, MA, United States).

### Isolation and purification of ginsentides

Ginsentide TP1, TP3 or TP8 were extracted from the dried flowers or seeds of *Panax* ginseng, *Panax quinquefolius,*and *Panax notoginseng* as previously described^18^. Briefly, the plant materials were pulverized and extracted with water. The extracts were filtered and subjected to flash chromatography using C18 powder (Grace Davison). 60% ethanol was used to elute the ginsentide-enriched fractions which were then loaded onto an SP Sepharose resin column (GE Healthcare, UK) and eluted with 1 M NaCl (pH 3.0). Further purification was done using preparative RP-HPLC (Shimadzu, Japan) with a C18 Grace Vydac column (250 × 22 mm) at a flow rate of 8 mL/min, linear gradient of 1%/min of 10–80% buffer B. Buffer A contained 0.05% (v/v) trifluoroacetic acid (TFA) in HPLC grade water, and buffer B contained 0.05% (v/v) TFA and 99.5% (v/v) acetonitrile (ACN). Resulting fractions were further purified using a semi-preparative C18 Vydac column (250 × 10 mm) at a flow rate of 3 mL/min with the same linear gradient. The identity of the ginsentide containing fractions were confirmed by matrix-assisted laser desorption/ionization-time of flight mass spectrometry (MALDI-TOF MS; AB SCIEX 5800 MALDI-TOF/TOF).

### Solid-phase peptide synthesis and oxidative folding

Synthetic ginsentide TP1 was synthesized by Fmoc-based solid-phase peptide synthesis on 2-Chlorotrityl chloride resin as previously described^16^. The linear precursor peptide was cleaved using a cocktail consisting of 92.5 % TFA, 2.5 % H_2_O, 2.5 % 1,2-ethanedithiol, and 2.5 % triisopropylsilane at room temperature for 30 min followed by precipitation with diethyl ether. The crude cleavage product was folded in 10% dimethyl sulfoxide (DMSO), 90% 0.1 M NH_4_HCO_3_ (pH 8), cystamine (10 equivalents), and cysteamine (100 equivalents) for overnight at 4 °C. Folded ginsentide TP1 was purified by preparative HPLC (250 x 21 mm, 5 μm; Phenomenex, USA). A linear gradient of mobile phase A (0.1 % TFA in H2O) and mobile phase B (0.1 % TFA in ACN) was used. The folded ginsentide TP1was identified using MALDI-TOF MS. The folding yield was approximately 30%. RP-HPLC and 2-dimensional-nuclear magnetic resonance (2D NMR) were performed to compare the physical properties of synthetic ginsentide TP1 to its native form (**Supplementary data S1 and S2**).

### Fluorescent labeling of ginsentide TP1

The lysine sidechain of native ginsentide TP1 was fluorescent-labeled using Cy3 NHS ester (Lumiprobe, USA) in 100 mM phosphate buffer (pH 7.8). Fluorescent labeling was carried out at room temperature for 16 h, and Cy3-TP1 was then identified and purified by RP-HPLC and MALDI-TOF MS.

### Immobilization of ginsentide TP1 onto agarose beads

Ginsentide TP1 was immobilized onto agarose beads using AminoLink^TM^ Immobilization Kit (Thermo Fisher Scientific, USA) according to manufacturer instructions.

### Confocal microscopy analysis

Live-cell confocal microscopy was conducted on HUVEC-CS (human umbilical vein endothelial) cells. HUVEC-CS were cultured in high glucose DMEM containing 1 mM sodium pyurvate, 4 mM L-glutamine supplemented with 10% fetal bovine serum (FBS) and 100 U/mL of penicillin and streptomycin in a 5% CO2 humidified incubator at 37 °C. To examine the intracellular distribution of Cy3-TP1, HUVEC-CS cells were seeded on an 8-well chamber slide (Ibidi, Germany). After incubation with Cy3-rT1, cells were stained with Hoechst 333241. The slides were imaged using a Zeiss LSM 710 confocal microscope.

### Cellular uptake analyses by flow cytometry

Cellular uptake of Cy3-TP1 was analyzed using flow cytometry, HUVEC-CS cells were incubated with Cy3-TP1 in serum-free medium at 37 °C. Following incubation, cells were harvested and collected by centrifugation at 500 g for 5 min. To quench extra-cellular fluorescence, cells were mixed with 150 µg/mL of trypan blue, and the samples were analyzed by flow cytometry. A total of 10,000 cells were analyzed using a BD LSR FortessaTM X-20 flow cytometer. For temperature-dependent uptake studies, HUVEC-CS cells were incubated at 4 °C for 30 min prior to incubation with Cy3-TP1 for 1 h at 4 °C. For endocytosis inhibitor studies, HUVEC-CS cells were pretreated with endocytosis inhibitors, including 50 µM dynasore, 50 µM ethylisopropylamiloride (EIPA), and 50 µg/mL nystatin for 30 min, followed by incubation with Cy3-TP1 for 1 h at 37 °C.

### Sample preparation and LC-MS/MS analysis

Affinity-enrichment mass spectrometry profiling was performed using immobilized ginsentide TP1 on agarose beads. After washing the beads with phosphate-buffered saline (PBS) three times, 600 μg of HUVEC-CS cell lysate was added to each tube and allowed to incubate overnight at 4 °C with gentle end-to-end rotation. After incubation, the resin was transferred to Pierce^®^spin columns and washed with PBS. 6X loading dye with 2-mercaptoethanol was added to the resin and heated for 10 min at 85 °C. The resultant mixture was centrifuged at 200 g for 1 min and resolved by electrophoresis at 100 V for 120 min using 15% SDS-PAGE. After the samples were resolved on the SDS-polyacrylamide gel, each sample lane was cut. The gel pieces were reduced with 10 mM DTT for 30 min at 60 °C and alkylated for 45 min with 55 mM iodoacetamide at room temperature. The samples were then subjected to in-gel tryptic digestion (Promega, USA) at 37 °C overnight. Tryptic peptides were extracted with 5% acetic acid in 50% ACN buffer and vacuum dried.

Tryptic peptides were then subjected to LC-MS/MS analysis using a Q Exactive mass spectrometer coupled with an online Dionex Ultimate 3000 RSLC nano-LC system (Thermo Fisher Scientific). Dried tryptic peptides were reconstructed in 0.1% formic acid solution and resolved using a Dionex EASY-spray column (PepMap C18; 3 μm; 100 Å) and injected through an EASY nanospray source. Data acquisition was performed using Xcaliber 2.2 (Thermo Scientific). Samples from experimental triplicate were run as technical duplicates. Mascot generic format files were generated from the raw data using Proteome Discoverer 1.4.1.14 software). In-house Mascot search engine (version 2.4.1; Matrix Science, UK) was used for the protein sequence database searches The UniProt Knowledgebase of human proteins (downloaded on Feb 19, 2016, including 140450 sequences and 47482854 residues) was used as a search database. Carbamidomethyl at cysteine was set as a static modification and methionine oxidation, asparagine and glutamine deamidation were used as dynamic modifications. Full trypsin digestion with maximum 2 missed cleavages was set as the digestion parameter. The peptide mass tolerance of ± 10 ppm (# ^13^C = 2) and a ± 0.02-Da for-fragment mass were set as mass tolerance parameters. The data with the significance threshold value of p<0.05 and “IgnoreIonsScoreBelow” value of 20 were extracted in *csv format. The False discovery rate (FDR) were calculated using target-decoy search strategy with cutoff set to ≤ 1%and proteins identified with multiple peptides were selected for the final analysis. Proteins identified in all three experimental replicates were only consider for final analysis. Online bioinformatics tool Reactome (https://reactome.org/) was used for the pathway analysis.

### Nitric oxide measurement

A specific fluorescence probe, 4, 5-diaminofluorescein diacetate (DAF-2 DA), was used to measure cellular NO. HUVEC-CS cells were seeded at a density of 1.0 x 105 cells/mL in a 96-well plate. Cells were first washed with Hank’s balanced salt solution (HBSS) prior to treatment with different concentrations of ginsentide TP1 (final DMSO concentration 0.1%) and 5 μM DAF-2 DA. To study the mechanisms involved, cells were pre-incubated with different inhibitors for 1 h followed by another 1 h treatment with ginsentide TP1. Treated cells were incubated for 1 h in the dark at 37 °C. Fluorescence intensity was measured at excitation wavelength 485 nm and emission wavelength 515 nm on an Infinite® 200 PRO series microplate reader. The results were presented as changes in normalized fluorescence intensity compared to control.

### Western blot analysis

HUVEC-CS cells treated with ginsentide TP1 were harvested with a scraper and lysed in CelLytic^TM^ M lysis buffer supplemented with protease inhibitor cocktail and phosphatase inhibitor cocktail on ice with frequent agitation for 30 min. The cell homogenates were centrifuged at 12000 rpm for 30 min at 4 °C and supernatants were collected. Protein concentrations were determined using bicinchoninic acid (BCA) reagent. Total protein (30 μg) mixed with SDS loading buffer was denatured at 70 °C for 10 min. Denatured protein samples were then electrophoretically resolved on SDS-PAGE and transferred onto a 0.2 μm PVDF membrane. The membranes were blocked for 1 h at room temperature with 5% BSA TBST. The membranes were incubated for 1 h at room temperature or overnight at 4 °C with rabbit primary antibody for total-akt (1:1000) (CAT No. 9272S, Cell Signaling Technology, USA) or rabbit phosphorylated akt (Ser 473) (1:1000) CAT No. 4060S, Cell Signaling Technology, USA) or rabbit phosphorylated eNOS (Ser 1177) (1:1000) (CAT No. 9570S, Cell Signaling Technology, USA) or mouse β-actin (1:10000) (CAT No. A5316, Merck, USA) diluted with 5% BSA in TBST. The membrane was then incubated for 1 h at room temperature with horseradish peroxidase-conjugated anti-rabbit (1:2000) (Cell Signaling Technology, USA) or anti-mouse secondary antibody (1:2000) (Cell Signaling Technology, USA) diluted with 5% BSA in TBST. The immune complexes were detected on X-ray film using Advansta chemiluminescent substrate. For re-probing, the membrane was stripped with guanidine hydrochloride stripping buffer (6 M guanidine hydrochloride, 0.2% Nonidet (NP-40), 0.1 M β-mercaptoethanol, 20 mM Tris–HCl, pH 7.5) twice for 5 min, followed by 5 min washing with TBST four times^59^. To determine the intensity of the protein targets, densitometric analysis was performed using NIH Image-J software (National Institutes of Health, USA) and was presented as relative to β-actin expression as an internal control.

### *Ex vivo* aortic ring contraction

All protocols and procedures were approved by the Animal Care and Welfare Committee of Institute of Materia Medica, Chinese Academy of Medical Sciences and Peking Union Medical College, Beijing, China. Aortic ring contraction studies were performed as previously described with slight modifications^60^. Following decapitation, the thoracic aorta was quickly removed, cleared of fat and connective tissue and cut into segments (rings of approximately 2 mm). Aortic rings were mounted in a 15 mL organ baths containing Krebs buffer (120.4 mM NaCl, 5.9 mM KCl, 2.5 mM CaCl_2_, 1.2 mM MgCl2, 1.2 mM NaH_2_PO_4_, 11.5 mM glucose, 25 mM NaHCO3, pH 7.5) at 37 °C with continuous 95% O_2_ and 5% CO_2_ supply. Endothelium-intact aortic rings were pre-constricted with 40 mM KCl, followed by a relaxant response to 30 mM acetylcholine to ensure their integrity and functionality. Aortic rings were then allowed to equilibrate for 40 min prior to isometric tension measurements of vascular contractility. The vascular responses of ginsentide TP1, TP3 and TP8 were characterized in 0.1 μM phenylephrine pre-contracted aortic rings. Isometric tension was measured by a force-displacement transducer and recorded using AcqKnowledge ACK100 Version 3.2 (BIOPAC Systems, USA). Aortic ring contractions were measured in volts and expressed as a percentage compared to control.

### Monocyte–endothelium adhesion assay

Monocyte-endothelium adhesion assay was performed as previously described. Briefly, HUVEC-CS cells were grown until confluent in a 96-black well clear bottom plate. The cells were pre-treated with different concentrations of ginsentide TP1 (1, 5, 10, 20 µM) or 100 µM GRGDSP or 20 µM of ginsenoside Rb1 or 20 µM of ginsenoside Rg1 or PBS for 30 min at 37 °C. After 30 min, HUVEC-CS cells were co-incubated with carboxyfluorescein succinimidyl ester (CFSE)-labeled THP-1 cells at a density of 1 × 10^6^ cells/well in serum-free RPMI medium for 1 h at 37 °C. Monocyte adhesion was measured by the fluorescence intensity of CFSE after washing five times with PBS using a fluorescence microplate reader.

### Pull-down assay

Pull-down assays was performed using immobilized ginsentide TP1 on agarose beads. Briefly, immobilized ginsentide TP1 was washed with PBS three times and incubated 600 μg of HUVEC-CS cell lysate was added to each tube and allowed to incubate overnight at 4 °C with gentle end-to-end rotation. After incubation, the mixture was transferred to Pierce®spin columns and washed 10 times with PBS. 6X loading dye with 2-mercaptoethanol was added to the resin and heated for 10 min at 85 °C. The resultant mixture was centrifuged at 200 g for 1 min and resolved using 8%-12% SDS-PAGE, at 100 V constant for 120 min. Blot transfer was performed onto a PVDF membrane at 250 mA for 120 min on ice. The blot was blocked with 5% BSA TBST before incubating overnight at 4 °C with anti-CD166 (CAT No. SC-74558, Santa Cruz Biotechnology, USA), or CD40 (CAT No. SC-13128, Santa Cruz Biotechnology, USA) anti-mouse antibody, or anti-ESAM (CAT No. SC-366580, Santa Cruz Biotechnology, USA) anti-rabbit antibody (1:500 in 5% BSA TBST). After incubation overnight, the membrane was washed with TBST at room temperature three times, 10 min each. The blot is then incubated with horseradish peroxidase-conjugated anti-mouse secondary antibody (1:2000) (Cell Signaling Technology, USA) or anti-rabbit (1:2000) (Cell Signaling Technology, USA), diluted with 5% BSA in TBST, for another 1 h at room temperature. The blot was washed five times 10 min each with TBST at room temperature before addition of chemiluminescence substrate and exposure on X-ray film.

### G-Protein Coupled Receptor profiling

G-Protein Coupled Receptor (GPCR) profiling was conducted using gpcrMAX^SM^ GPCR Assay Panel, a functional cell-based assay, by DiscoverX, USA as a service contract. The effects of ginsentide TP1 on 168 GPCRs were screened in both agonist and antagonist modes.

### P2Y12 **β**-Arrestin recruitment assay

P2Y12 β-Arrestin recruitment assay was conducted as previous described by DiscoverX (USA) as a service contract. The effects of ginsentide TP1, ginsenoside Rb1, or ginsenoside Rg1 on ADP-induced P2Y12 activation were examined in antagonist modes.

### *Ex vivo* ADP-induced platelet aggregation assay

*Ex vivo* ADP-induced platelet aggregation assay was performed as previously described^61^. Sprague-Dawley (SD) rats (300–350 g body weight) obtained from the Animal Center of Chinese Academy of Medical Sciences were kept in a temperature-controlled environment (24 °C) under a 12/12-h light/dark cycle, fed with normal rat chow and water ad libitum. All protocols and procedures were approved by the Animal Care and Welfare Committee of Institute of Materia Medica, Chinese Academy of Medical Sciences and Peking Union Medical College, Beijing, China. Intravenous injection of ginsentide TP1 (10 mg/kg) or oral gavage of ginsentide TP1 (3, 10, 30 mg/kg) to SD rats were conducted once daily for 3 days. The control rats were given 0.9 % normal saline. All *ex vivo* experiments were conducted 30 min after the final pre-treatment. Rats were anesthetized with 10 % chloral hydrate (3 mL/kg, i.p.), and blood was collected from the abdominal aorta and anticoagulated with sodium citrate (3.8 %; 1:9, *v*/*v*). Platelet-rich plasma (PRP) was obtained by centrifuging blood at 1000 rpm for 10 min at room temperature. And, the remainder further centrifuged at 3000 rpm for 10 min to obtain platelet-poor plasma (PPP). The platelet counts of PRP were adjusted to 4C×C10^8^/mL with PPP. Platelet aggregation was determined by Born’s method (Born GV (1962) Aggregation of blood platelets by adenosine diphosphate and its reversal. Nature 194:927–929) using a four-channel aggregometer (LBY-NJ4, Beijing Precil Instrument Co.Ltd., China). 300 μL PRP was placed in a cuvette and stirred with rotor at 37 °C for 5 min. ADP (2.5 or 5 μM) was added to initiate platelet aggregation. Platelet aggregation was determined by calibrating the equipment at 0 % light transmission for PRP and at 100 % for PPP. All experiments were conducted within 2 h of blood collection. The percentage of platelet aggregation was determined by optical density. The aggregation curve was recorded for 5 min, and the peak value was received and served as the maximal aggregation (MAG). Inhibition% of platelet aggregation was calculated from the formula: inhibition%C=C(MAG of controlC−CMAG of tests) / MAG of controlC×C100 %.

### *In vivo* acute pulmonary thrombosis

The anti-thrombotic effects of ginsentide TP1 was studied using an *in vivo* acute thrombotic mouse model^62^ conducted by Thousand Dimensions (Beijing) Science and Technology Co., Ltd, Beijing, China, as a service contract. Male Institute of Cancer Research (ICR) mice ( 23-25 g body weight) were purchased from Beijing HFK bioscience CO.,LTD, Beijing, China. Briefly, Animals were randomly divided into 6 groups: sham group (N=5), control group (N=8), clopidogrel group (50 mg/kg) (N=7), aspirin group (100 mg/kg) (N=9), ginsentide TP1 group (10 mg/kg) (N=7), and ginsentide TP1 group (30 mg/kg) (N=7). Animals in each group were given the corresponding dose once a day at a dose volume of 10 mL/kg for 3 consecutive days, and the control group was given the corresponding volume of distilled water. Animals received a saline injection containing rat tail collagen (1.5 mg/kg) and epinephrine (0.5 mg/kg) via tail vein 1 h after dosing on Day 3. After intravenous injection for 15 s, the death of animals were recorded. 30 min after tail intravenous injection, the animals were sacrificed and their lungs were dissected. The dissected lungs were washed with saline, dried with filter paper, and weighed. In the sham group, tail vein injection was not performed and the animals were sacrificed three days later for lung measurement. The statistical indicators were lung coefficient and mortality. Lung coefficient= Lung tissue weight/Body weight*100%

### Forced swimming test

Male ICR mice (220-240 mg body weight) obtained from the Animal Center of Chinese Academy of Medical Sciences were kept in a temperature-controlled environment (24 °C) under a 12/12-h light/dark cycle, with food and water available. The animals were given 1 week to acclimatize to the housing conditions before the beginning of the experiments. All protocols and procedures were approved by the Animal Care and Welfare Committee of Institute of Materia Medica, Chinese Academy of Medical Sciences and Peking Union Medical College, Beijing, China. The Forced Swimming Test (FST) was performed as previously described^63, 64^. The animals were divided into control and three experimental groups as follows: vehicle-control (0.9% physiological saline) (N=14), fluoxetine (20 mg/kg) (N=13), ginsentide TP1 (10 mg/kg) (N=6), ginsentide TP1 (30 mg/kg) (N=13). All treatments were administered by oral (p.o.) gavage in a volume of 10 mL/kg body weight. Fluoxetine and ginsentide TP1 were administered to mice 30 min and 60 min before and after the FST, respectively. Mice were individually placed in a glass cylinder (20 cm in height, 14 cm in diameter) filled 10 cm high with water (25 ± 2 °C). All animals were forced to swim for 6 min, and the duration of immobility was observed and measured during the final 4 min interval of the test. Immobility period was regarded as the time spent by the mouse floating in the water without struggling and making only those movements necessary to keep its head above the water. The test sessions were recorded by a video camera and scored by an observer blind to treatment.

### Tail suspension test

Male ICR mice (220-240 mg body weight) obtained from the Animal Center of Chinese Academy of Medical Sciences were kept in a temperature-controlled environment (24°C) under a 12/12-h light/dark cycle, with food and water available. The animals were given 1 week to acclimatize to the housing conditions before the beginning of the experiments. All protocols and procedures were approved by the Animal Care and Welfare Committee of Institute of Materia Medica, Chinese Academy of Medical Sciences and Peking Union Medical College, Beijing, China. The Tail Suspension Test (TST) was conducted as previously described^52^. The animals were divided into control and three experimental groups as follows: vehicle-control (0.9% physiological saline) (N=13), fluoxetine group (20 mg/kg) (N=13), ginsentide TP1 group (10 mg/kg) (N=20), ginsentide TP1 group (30 mg/kg) (N=12). All treatments were administered by oral (p.o.) gavage in a volume of 10 mL/kg body weight. Fluoxetine and ginsentide TP1 were administered to mice a day before as well as 30 min and 60 min, respectively, before the TST. Briefly, mice were individually suspended by tail with a clamp (1 cm from the tip of the end) in a box (25 × 25 × 30 cm) with the head 5 cm from the bottom. Testing was carried out in a darkened room with minimal background noise. A mouse was suspended for a total of 6 min, and the duration of immobility was recorded during the final 4 min interval of the test. Mice were considered immobile only when they hung passively and completely motionless. The test sessions were recorded by a video camera and scored by an observer blind to treatment.

### Statistics

Statistical comparisons were performed using GraphPad Version 8.2.1 (US). Data were analyzed using one-way analysis of variance (ANOVA) followed by Newman-Keuls *post hoc* tests. Data were expressed as mean±S.D. and *p*<0.05 was considered to be statistically significant.

### Data Availability Section

The affinity-enriched mass spectrometry data and Mascot search files generated in this study has been deposited in the ProteomeXchange Consortium (http://proteomecentral.proteomexchange.org) via the jPOSTrepo (Japan ProteOme STandard Repository) partner repository^65^ under the accession codes of PXD039905 and JPST001994, respectively.

## Author contributions

S.L. and A.K. performed the experiments and wrote the manuscript. B.D. collected and analyzed the proteomic profiling and target identification data. X.Z. performed the chemical synthesis. N.F. performed the animal studies. S.K.S. oversaw the analysis of the proteomics analysis. C.F.L. oversaw the chemical synthesis. X.W. oversaw the animal studies. J.P.T. conceived the idea and edited the manuscript. All authors have read and approved the manuscript.

## Competing interests

The authors declare that the research was conducted in the absence of any commercial or financial relationships that could be construed as a potential conflict of interest.

## Acknowledgements

This research was supported in part by the Nanyang Technological University Internal Funding -Synzyme and Natural Products (SYNC) and the AcRF Tier 3 funding (MOE2016-T3-1-003).

## Notes

http://proteomecentral.proteomexchange.org

### Competing Interest Statement

The authors have declared no competing interest.

### Summary of Updates

This version of the manuscript was revised to update the funding source.

