## Supplementary material for "Decoding the Cure-all Effects of Ginseng": Dataset 1

**Dataset 1: List of proteins pulled-down by ginsentide TP1**

| **#** | **Accession** | **Gene** | **Protein** |
| --- | --- | --- | --- |
| 1 | A0A087WX41 | CLTCL1 | Clathrin heavy chain 2 |
| 2 | M0R1I1 | TUBB4A | Tubulin beta-4A chain |
| 3 | A0A0U1RQF0 | FASN | Fatty acid synthase |
| 4 | Q15738 | NSDHL | Sterol-4-alpha-carboxylate 3-dehydrogenase, decarboxylating |
| 5 | P33176 | KIF5B | Kinesin-1 heavy chain |
| 6 | C9JPC0 | PSMD2 | 26S proteasome non-ATPase regulatory subunit 2 |
| 7 | Q9Y490 | TLN1 | Talin-1 |
| 8 | Q9Y6G9 | DYNC1LI1 | Cytoplasmic dynein 1 light intermediate chain 1 |
| 9 | X6RJ95 | PTGES2 | Prostaglandin E synthase 2 |
| 10 | A0A087WXS7 | ASNA1 | ATPase ASNA1 |
| 11 | D6RBV2 | LMAN2 | Vesicular integral-membrane protein VIP36 |
| 12 | Q9Y394 | DHRS7 | Dehydrogenase/reductase SDR family member 7 |
| 13 | O00487 | PSMD14 | 26S proteasome non-ATPase regulatory subunit 14 |
| 14 | G3V542 | TUBB3 | Tubulin beta-3 chain |
| 15 | A2IDC6 | MRPL28 | 39S ribosomal protein L28, mitochondrial |
| 16 | Q9UNM6 | PSMD13 | 26S proteasome non-ATPase regulatory subunit 13 |
| 17 | H7C2Q3 | PSMD2 | 26S proteasome non-ATPase regulatory subunit 2 |
| 18 | P10301 | RRAS | Ras-related protein R-Ras |
| 19 | Q9P015 | MRPL15 | 39S ribosomal protein L15, mitochondrial |
| 20 | P51665 | PSMD7 | 26S proteasome non-ATPase regulatory subunit 7 |
| 21 | P09132 | SRP19 | Signal recognition particle 19 kDa protein |
| 22 | F5H6X6 | GANAB | Neutral alpha-glucosidase AB |
| 23 | P63244 | GNB2L1 | Guanine nucleotide-binding protein subunit beta-2-like 1 |
| 24 | A0A0D9SF54 | SPTAN1 | Spectrin alpha chain, non-erythrocytic 1 |
| 25 | Q99614 | TTC1 | Tetratricopeptide repeat protein 1 |
| 26 | P61981 | YWHAG | 14-3-3 protein gamma |
| 27 | P20020 | ATP2B1 | Plasma membrane calcium-transporting ATPase 1 |
| 28 | Q6YN16 | HSDL2 | Hydroxysteroid dehydrogenase-like protein 2 |
| 29 | Q6IAA8 | LAMTOR1 | Ragulator complex protein LAMTOR1 |
| 30 | K7EJE8 | LONP1 | Lon protease homolog |
| 31 | J3KN29 | PSMD9 | 26S proteasome non-ATPase regulatory subunit 9 |
| 32 | A0A087X1Z3 | PSME2 | Proteasome activator complex subunit 2 |
| 33 | H0YEB6 | SSSCA1 | Sjoegren syndrome/scleroderma autoantigen 1 |
| 34 | Q7L1Q6 | BZW1 | Basic leucine zipper and W2 domain-containing protein 1 |
| 35 | P51398 | DAP3 | 28S ribosomal protein S29, mitochondrial |
| 36 | H0YFX9 | H2AFJ | Histone H2A |
| 37 | A0A0A0MQS1 | PYCRL | Pyrroline-5-carboxylate reductase |
| 38 | Q9BRP8 | PYM1 | Partner of Y14 and mago |
| 39 | P26640 | VARS | Valine--tRNA ligase |
| 40 | P61163 | ACTR1A | Alpha-centractin |
| 41 | H7C543 | ALCAM | CD166 antigen |
| 42 | C9J8P9 | CLTA | Clathrin light chain A |
| 43 | Q96CS3 | FAF2 | FAS-associated factor 2 |
| 44 | I3L0E3 | hCG-1984214 | HCG1984214, isoform CRA_a |
| 45 | Q58FF6 | HSP90AB4P | Putative heat shock protein HSP 90-beta 4 |
| 46 | P08648 | ITGA5 | Integrin alpha-5 |
| 47 | P33993 | MCM7 | DNA replication licensing factor MCM7 |
| 48 | X6RAY8 | MRPL4 | 39S ribosomal protein L4, mitochondrial |
| 49 | P82912 | MRPS11 | 28S ribosomal protein S11, mitochondrial |
| 50 | Q92665 | MRPS31 | 28S ribosomal protein S31, mitochondrial |
| 51 | O15031 | PLXNB2 | Plexin-B2 |
| 52 | O00592 | PODXL | Podocalyxin |
| 53 | H0YJX2 | PSMC6 | 26S protease regulatory subunit 10B |
| 54 | Q9BTV4 | TMEM43 | Transmembrane protein 43 |
| 55 | Q9NRR5 | UBQLN4 | Ubiquilin-4 |
| 56 | P62258 | YWHAE | 14-3-3 protein epsilon |
| 57 | Q16543 | CDC37 | Hsp90 co-chaperone Cdc37 |
| 58 | P19022 | CDH2 | Cadherin-2 |
| 59 | M0R208 | CLPP | ATP-dependent Clp protease proteolytic subunit |
| 60 | F8VVA7 | COPZ1 | Coatomer subunit zeta-1 |
| 61 | A0A0C4DGQ8 | DHRS7B | Dehydrogenase/reductase SDR family member 7B |
| 62 | E7EUU4 | EIF4G1 | Eukaryotic translation initiation factor 4 gamma 1 |
| 63 | Q13283 | G3BP1 | Ras GTPase-activating protein-binding protein 1 |
| 64 | E9PKU7 | GANAB | Neutral alpha-glucosidase AB |
| 65 | P30499 | HLA-C | HLA class I histocompatibility antigen, Cw-1 alpha chain |
| 66 | P43246 | MSH2 | DNA mismatch repair protein Msh2 |
| 67 | Q8TC12 | RDH11 | Retinol dehydrogenase 11 |
| 68 | B1AJQ6 | STX12 | Syntaxin-12 |
| 69 | Q5T8U5 | SURF4 | Surfeit 4 |
| 70 | Q9NYU2 | UGGT1 | UDP-glucose:glycoprotein glucosyltransferase 1 |
| **#** | **Accession** | **Gene** | **Protein** |
| 71 | E9PG15 | YWHAQ | 14-3-3 protein theta |
| 72 | E9PJN0 | ACOT8 | Acyl-coenzyme A thioesterase 8 |
| 73 | P42025 | ACTR1B | Beta-centractin |
| 74 | E9PG40 | APP | Amyloid beta A4 protein |
| 75 | C9JMD7 | BCAP31 | B-cell receptor-associated protein 31 |
| 76 | P09497 | CLTB | Clathrin light chain B |
| 77 | J3KS22 | DCXR | L-xylulose reductase |
| 78 | D6RBD7 | EEF1E1 | Eukaryotic translation elongation factor 1 epsilon-1 |
| 79 | Q14152 | EIF3A | Eukaryotic translation initiation factor 3 subunit A |
| 80 | Q9NUQ9 | FAM49B | Protein FAM49B |
| 81 | J3QKQ5 | KPNB1 | Importin subunit beta-1 |
| 82 | P08582 | MFI2 | Melanotransferrin |
| 83 | P82675 | MRPS5 | 28S ribosomal protein S5, mitochondrial |
| 84 | P52701 | MSH6 | DNA mismatch repair protein Msh6 |
| 85 | P61601 | NCALD | Neurocalcin-delta |
| 86 | B3KQV6 | PPP2R1A | Serine/threonine-protein phosphatase 2A 65 kDa regulatory subunit A alpha |
| 87 | Q5BJH1 | PSAP | PSAP protein |
| 88 | O14828 | SCAMP3 | Secretory carrier-associated membrane protein 3 |
| 89 | C9J0K6 | SRI | Sorcin |
| 90 | H3BS86 | STUB1 | E3 ubiquitin-protein ligase CHIP |
| 91 | D6RC71 | STX18 | Syntaxin-18 |
| 92 | J3KQ45 | TGOLN2 | Trans-Golgi network integral membrane protein 2 |
| 93 | O94826 | TOMM70A | Mitochondrial import receptor subunit TOM70 |
| 94 | Q9UMX0 | UBQLN1 | Ubiquilin-1 |
| 95 | E7ETF4 | UGDH | UDP-glucose 6-dehydrogenase |
| 96 | Q9Y6D5 | ARFGEF2 | Brefeldin A-inhibited guanine nucleotide-exchange protein 2 |
| 97 | H7C547 | ATP1B3 | Sodium/potassium-transporting ATPase subunit beta-3 |
| 98 | P32970 | CD70 | CD70 antigen |
| 99 | P08574 | CYC1 | Cytochrome c1, heme protein, mitochondrial |
| 100 | Q96PD2 | DCBLD2 | Discoidin, CUB and LCCL domain-containing protein 2 |
| 101 | C9JYS6 | EGFR | Epidermal growth factor receptor |
| 102 | H0YDT6 | EIF3F | Eukaryotic translation initiation factor 3 subunit F |
| 103 | V9GZ76 | EPRS | Bifunctional glutamate/proline--tRNA ligase |
| 104 | Q96AP7 | ESAM | Endothelial cell-selective adhesion molecule |
| 105 | Q9Y3D0 | FAM96B | Mitotic spindle-associated MMXD complex subunit MIP18 |
| 106 | Q5T196 | FLAD1 | FAD synthase |
| 107 | P51991 | HNRNPA3 | Heterogeneous nuclear ribonucleoprotein A3 |
| 108 | P14866 | HNRNPL | Heterogeneous nuclear ribonucleoprotein L |
| 109 | P17301 | ITGA2 | Integrin alpha-2 |
| 110 | V9GYZ1 | ITGB5 | Integrin beta |
| 111 | H7C3W8 | LRPPRC | Leucine-rich PPR motif-containing protein, mitochondrial |
| 112 | Q9H9J2 | MRPL44 | 39S ribosomal protein L44, mitochondrial |
| 113 | Q00653 | NFKB2 | Nuclear factor NF-kappa-B p100 subunit |
| 114 | Q6DKJ4 | NXN | Nucleoredoxin |
| 115 | A0A0A0MSW4 | PITPNB | Phosphatidylinositol transfer protein beta isoform |
| 116 | P55036 | PSMD4 | 26S proteasome non-ATPase regulatory subunit 4 |
| 117 | R4GMR5 | PSMD8 | 26S proteasome non-ATPase regulatory subunit 8 |
| 118 | A8MXQ1 | PTTG1IP | Pituitary tumor-transforming gene 1 protein-interacting protein |
| 119 | Q9GZT4 | SRR | Serine racemase |
| 120 | P31948 | STIP1 | Stress-induced-phosphoprotein 1 |
| 121 | C9JA93 | TBC1D15 | TBC1 domain family member 15 |
| 122 | K7ERV3 | TK1 | Thymidine kinase |
| 123 | A0A0A0MSK5 | TOR1AIP1 | Torsin-1A-interacting protein 1 |
| 124 | Q5HY81 | UBL4A | Ubiquitin-like protein 4A |
| 125 | P31946 | YWHAB | 14-3-3 protein beta/alpha |
| 126 | O95159 | ZFPL1 | Zinc finger protein-like 1 |
| 127 | P49189 | ALDH9A1 | 4-trimethylaminobutyraldehyde dehydrogenase |
| 128 | O75915 | ARL6IP5 | PRA1 family protein 3 |
| 129 | J3QSY6 | ATP2A2 | Sarcoplasmic/endoplasmic reticulum calcium ATPase 2 |
| 130 | G3V325 | ATP5J2-PTCD1 | Protein ATP5J2-PTCD1 |
| 131 | Q96BR5 | COA7 | Cytochrome c oxidase assembly factor 7 |
| 132 | P26232 | CTNNA2 | Catenin alpha-2 |
| 133 | Q16531 | DDB1 | DNA damage-binding protein 1 |
| 134 | B8ZZ50 | EIF4E2 | Eukaryotic translation initiation factor 4E type 2 |
| 135 | Q8TAE8 | GADD45GIP1 | Growth arrest and DNA damage-inducible proteins-interacting protein 1 |
| 136 | B8ZZA8 | GLS | Glutaminase kidney isoform, mitochondrial |
| 137 | Q9NQX3 | GPHN | Gephyrin |
| 138 | P35914 | HMGCL | Hydroxymethylglutaryl-CoA lyase, mitochondrial |
| 139 | P32004 | L1CAM | Neural cell adhesion molecule L1 |
| 140 | H0Y9G6 | MRPL3 | 39S ribosomal protein L3, mitochondrial |
| 141 | C9JG87 | MRPL39 | 39S ribosomal protein L39, mitochondrial |
| 142 | H7C5L9 | MRPS22 | 28S ribosomal protein S22, mitochondrial |
| **#** | **Accession** | **Gene** | **Protein** |
| 143 | H7C3G9 | NAGK | N-acetyl-D-glucosamine kinase |
| 144 | A0A087WZQ7 | NAPB | Beta-soluble NSF attachment protein |
| 145 | A0A0A0MRJ6 | PCMT1 | Protein-L-isoaspartate O-methyltransferase |
| 146 | C9J2Y9 | POLR2B | DNA-directed RNA polymerase |
| 147 | B1AJY7 | PSMD10 | 26S proteasome non-ATPase regulatory subunit 10 |
| 148 | P11216 | PYGB | Glycogen phosphorylase, brain form |
| 149 | E7ES60 | RAB34 | Ras-related protein Rab-34, isoform NARR |
| 150 | B4E0Y9 | STK26 | Serine/threonine-protein kinase 26 |
| 151 | H3BT89 | TMX3 | Protein disulfide-isomerase TMX3 |
| 152 | Q8NFQ8 | TOR1AIP2 | Torsin-1A-interacting protein 2 |
| 153 | Q9UHD9 | UBQLN2 | Ubiquilin-2 |
| 154 | Q04917 | YWHAH | 14-3-3 protein eta |
| 155 | Q16186 | ADRM1 | Proteasomal ubiquitin receptor ADRM1 |
| 156 | H3BQB1 | APRT | Adenine phosphoribosyltransferase |
| 157 | Q5QP56 | BCL2L1 | Bcl-2-like protein 1 |
| 158 | Q02338 | BDH1 | D-beta-hydroxybutyrate dehydrogenase, mitochondrial |
| 159 | I3L3B0 | C1QBP | Complement component 1 Q subcomponent-binding protein, mitochondrial |
| 160 | Q96S66 | CLCC1 | Chloride channel CLIC-like protein 1 |
| 161 | J3KRF5 | CLTC | Clathrin heavy chain 1 |
| 162 | C9J384 | CMSS1 | Protein CMSS1 |
| 163 | Q5T063 | COQ3 | Ubiquinone biosynthesis O-methyltransferase, mitochondrial |
| 164 | Q9UFM8 | DKFZp566H1924 | Neuroplastin |
| 165 | H0Y368 | DPM1 | Dolichol-phosphate mannosyltransferase subunit 1 |
| 166 | E9PHZ1 | ECE1 | Endothelin-converting enzyme 1 |
| 167 | P26641 | EEF1G | Elongation factor 1-gamma |
| 168 | A0A087WZK9 | EIF3H | Eukaryotic translation initiation factor 3 subunit H |
| 169 | E9PEB5 | FUBP1 | Far upstream element-binding protein 1 |
| 170 | O75600 | GCAT | 2-amino-3-ketobutyrate coenzyme A ligase, mitochondrial |
| 171 | Q6Y7W6 | GIGYF2 | PERQ amino acid-rich with GYF domain-containing protein 2 |
| 172 | F8W026 | HSP90B1 | Endoplasmin |
| 173 | H0YBG6 | HSPA9 | Stress-70 protein, mitochondrial |
| 174 | Q9NSE4 | IARS2 | Isoleucine--tRNA ligase, mitochondrial |
| 175 | A0A0C4DFN3 | MGLL | Monoglyceride lipase |
| 176 | Q9UBG0 | MRC2 | C-type mannose receptor 2 |
| 177 | A6NJD9 | MRPL23 | 39S ribosomal protein L23, mitochondrial |
| 178 | A0A087WU62 | MRPL45 | 39S ribosomal protein L45, mitochondrial |
| 179 | P82914 | MRPS15 | 28S ribosomal protein S15, mitochondrial |
| 180 | A0A087WZG2 | MST1R | Macrophage-stimulating protein receptor |
| 181 | Q9Y697 | NFS1 | Cysteine desulfurase, mitochondrial |
| 182 | Q9UKX7 | NUP50 | Nuclear pore complex protein Nup50 |
| 183 | H3BMX0 | NUP93 | Nuclear pore complex protein Nup93 |
| 184 | H0YAS6 | PABPC1 | Polyadenylate-binding protein 1 |
| 185 | H3BSP4 | PCBP2 | Poly(rC)-binding protein 2 |
| 186 | P67775 | PPP2CA | Serine/threonine-protein phosphatase 2A catalytic subunit alpha isoform |
| 187 | P25788 | PSMA3 | Proteasome subunit alpha type-3 |
| 188 | Q06323 | PSME1 | Proteasome activator complex subunit 1 |
| 189 | Q92692 | PVRL2 | Nectin-2 |
| 190 | P54727 | RAD23B | UV excision repair protein RAD23 homolog B |
| 191 | K7EIY6 | RNF126 | E3 ubiquitin-protein ligase RNF126 |
| 192 | Q99942 | RNF5 | E3 ubiquitin-protein ligase RNF5 |
| 193 | Q9Y6Y8 | SEC23IP | SEC23-interacting protein |
| 194 | A0A0G2JRX5 | SMN1 | Survival motor neuron protein |
| 195 | A0A087WUZ3 | SPTBN1 | Spectrin beta chain, non-erythrocytic 1 |
| 196 | E9PFW8 | SQSTM1 | Sequestosome-1 |
| 197 | K7EQB1 | STX8 | Syntaxin-8 |
| 198 | Q8TC07 | TBC1D15 | TBC1 domain family member 15 |
| 199 | Q96JH7 | VCPIP1 | Deubiquitinating protein VCIP135 |
| 200 | Q9HAV4 | XPO5 | Exportin-5 |
| 201 | Q92667 | AKAP1 | A-kinase anchor protein 1, mitochondrial |
| 202 | G3V1C3 | API5 | Apoptosis inhibitor 5 |
| 203 | B1ALK7 | ARHGEF7 | Rho guanine nucleotide exchange factor 7 |
| 204 | A0A0U1RQU3 | ATP2B1 | Plasma membrane calcium-transporting ATPase 1 |
| 205 | H0Y7S3 | ATP2B2 | Plasma membrane calcium-transporting ATPase 2 |
| 206 | A0A087X1D0 | CD40 | CD40 antigen (TNF receptor superfamily member 5), isoform CRA_c |
| 207 | F8W7Q4 | FAM162A | Protein FAM162A |
| 208 | P00367 | GLUD1 | Glutamate dehydrogenase 1, mitochondrial |
| 209 | F5GYK7 | GPD2 | Glycerol-3-phosphate dehydrogenase |
| 210 | D6RCN7 | IPO11 | Importin-11 |
| 211 | O95373 | IPO7 | Importin-7 |
| 212 | J3KS65 | KPNA2 | Importin subunit alpha-1 |
| 213 | X6RLN4 | LARP4 | La-related protein 4 |
| 214 | G3V1P5 | MED15 | Mediator of RNA polymerase II transcription subunit 15 |
| **#** | **Accession** | **Gene** | **Protein** |
| 215 | P08473 | MME | Neprilysin |
| 216 | Q9H2W6 | MRPL46 | 39S ribosomal protein L46, mitochondrial |
| 217 | H3BM93 | NUP93 | Nuclear pore complex protein Nup93 |
| 218 | G3V5L5 | PRMT5 | Protein arginine N-methyltransferase 5 |
| 219 | A0A0U1RQQ4 | PROCR | Endothelial protein C receptor |
| 220 | Q4VXH0 | PSMD5 | 26S proteasome non-ATPase regulatory subunit 5 |
| 221 | P23468 | PTPRD | Receptor-type tyrosine-protein phosphatase delta |
| 222 | E9PLA6 | SERPINH1 | Serpin H1 |
| 223 | D6RFW1 | SGTB | Small glutamine-rich tetratricopeptide repeat-containing protein beta |
| 224 | Q99808 | SLC29A1 | Equilibrative nucleoside transporter 1 |
| 225 | J3KRY3 | SNRPN | Small nuclear ribonucleoprotein-associated protein N |
| 226 | A0A087WVA8 | TEX2 | Testis-expressed sequence 2 protein |
| 227 | A0A087WWM0 | TRAPPC3 | Trafficking protein particle complex subunit 3 |
| 228 | Q6RW13 | AGTRAP | Type-1 angiotensin II receptor-associated protein |
| 229 | B4DIW9 | ANKRD28 | Serine/threonine-protein phosphatase 6 regulatory ankyrin repeat subunit A |
| 230 | Q8N2F6 | ARMC10 | Armadillo repeat-containing protein 10 |
| 231 | F8VZ13 | CDK4 | Cyclin-dependent kinase 4 |
| 232 | E7EMJ5 | CTNNB1 | Catenin beta-1 |
| 233 | A0A0G2JH68 | DIAPH1 | Protein diaphanous homolog 1 |
| 234 | Q14213 | EBI3 | Interleukin-27 subunit beta |
| 235 | I3L2C7 | GEMIN4 | Gem-associated protein 4 |
| 236 | G3V1U5 | GOLT1B | Golgi transport 1 homolog B (S. cerevisiae), isoform CRA_c |
| 237 | H0YAA0 | H0YAA0 | Uncharacterized protein |
| 238 | A8MUF7 | HBE1 | Hemoglobin subunit epsilon |
| 239 | H0YA49 | ITGA3 | Integrin alpha-3 |
| 240 | P13473 | LAMP2 (CD107b) | Lysosome-associated membrane glycoprotein 2 |
| 241 | Q9BYD1 | MRPL13 | 39S ribosomal protein L13, mitochondrial |
| 242 | Q8N5N7 | MRPL50 | 39S ribosomal protein L50, mitochondrial |
| 243 | P82664 | MRPS10 | 28S ribosomal protein S10, mitochondrial |
| 244 | E5RJ73 | MRPS27 | 28S ribosomal protein S27, mitochondrial |
| 245 | Q9UBV8 | PEF1 | Peflin |
| 246 | Q9BTU6 | PI4K2A | Phosphatidylinositol 4-kinase type 2-alpha |
| 247 | J3KSK1 | PSMD12 | 26S proteasome non-ATPase regulatory subunit 12 |
| 248 | Q13308 | PTK7 | Inactive tyrosine-protein kinase 7 |
| 249 | F2Z2V6 | QARS | Glutamine--tRNA ligase |
| 250 | R4GMU7 | RPL7L1 | 60S ribosomal protein L7-like 1 |
| 251 | J3KTD2 | RTTN | Rotatin |
| 252 | H0Y362 | SLC30A7 | Zinc transporter 7 |
| 253 | Q0IIM8 | TBC1D8B | TBC1 domain family member 8B |
| 254 | Q8WUY1 | THEM6 | Protein THEM6 |
| 255 | H0Y4W2 | TRRAP | Transformation/transcription domain-associated protein |
| 256 | K7EKG2 | TXNL1 | Thioredoxin-like protein 1 |
