## Supplementary data S1-S8 for "Decoding the Cure-all Effects of Ginseng"

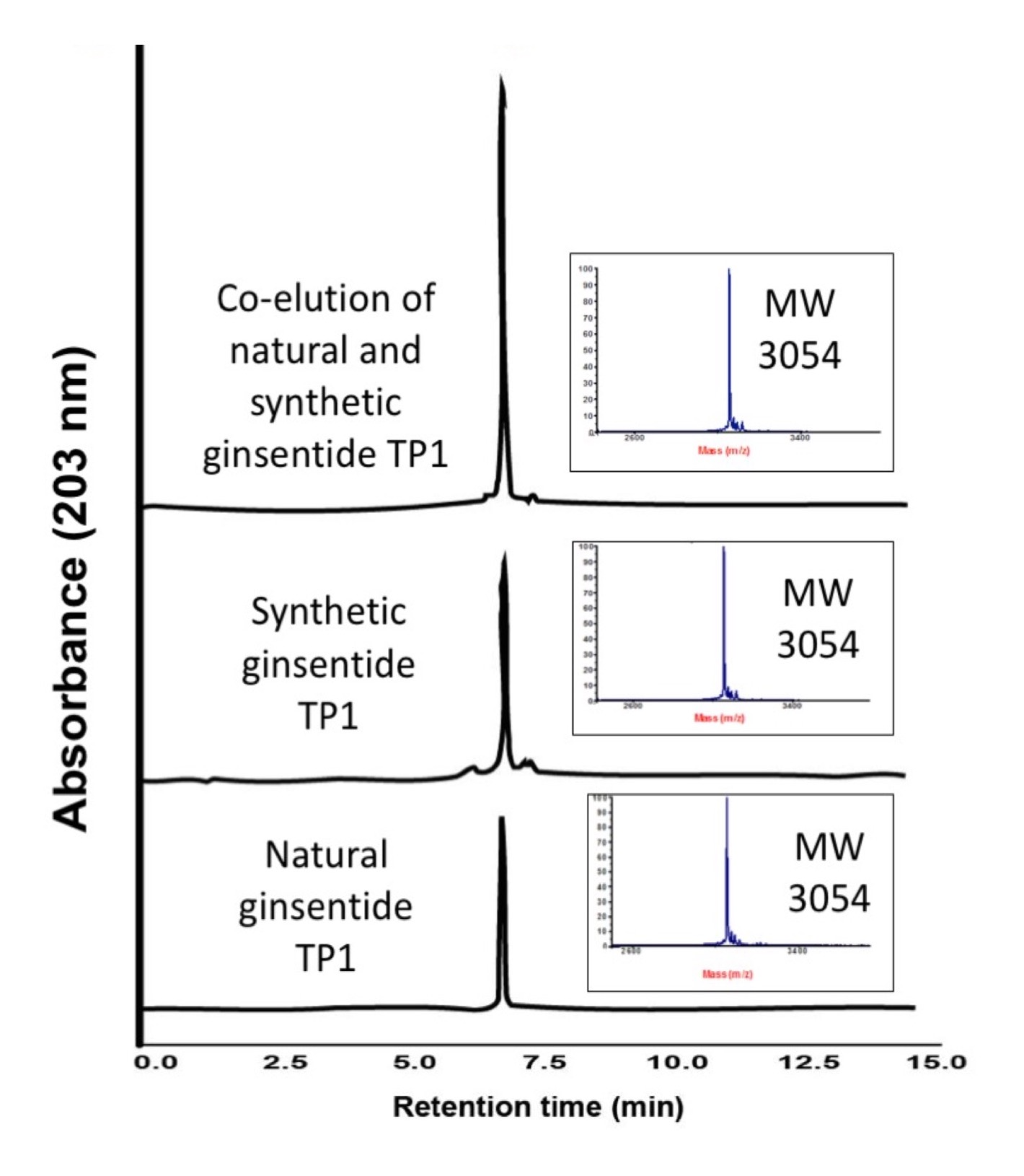

**Supplementary data S1.** Co-elution of natural and synthetic ginsentide TP1 using RP-HPLC.

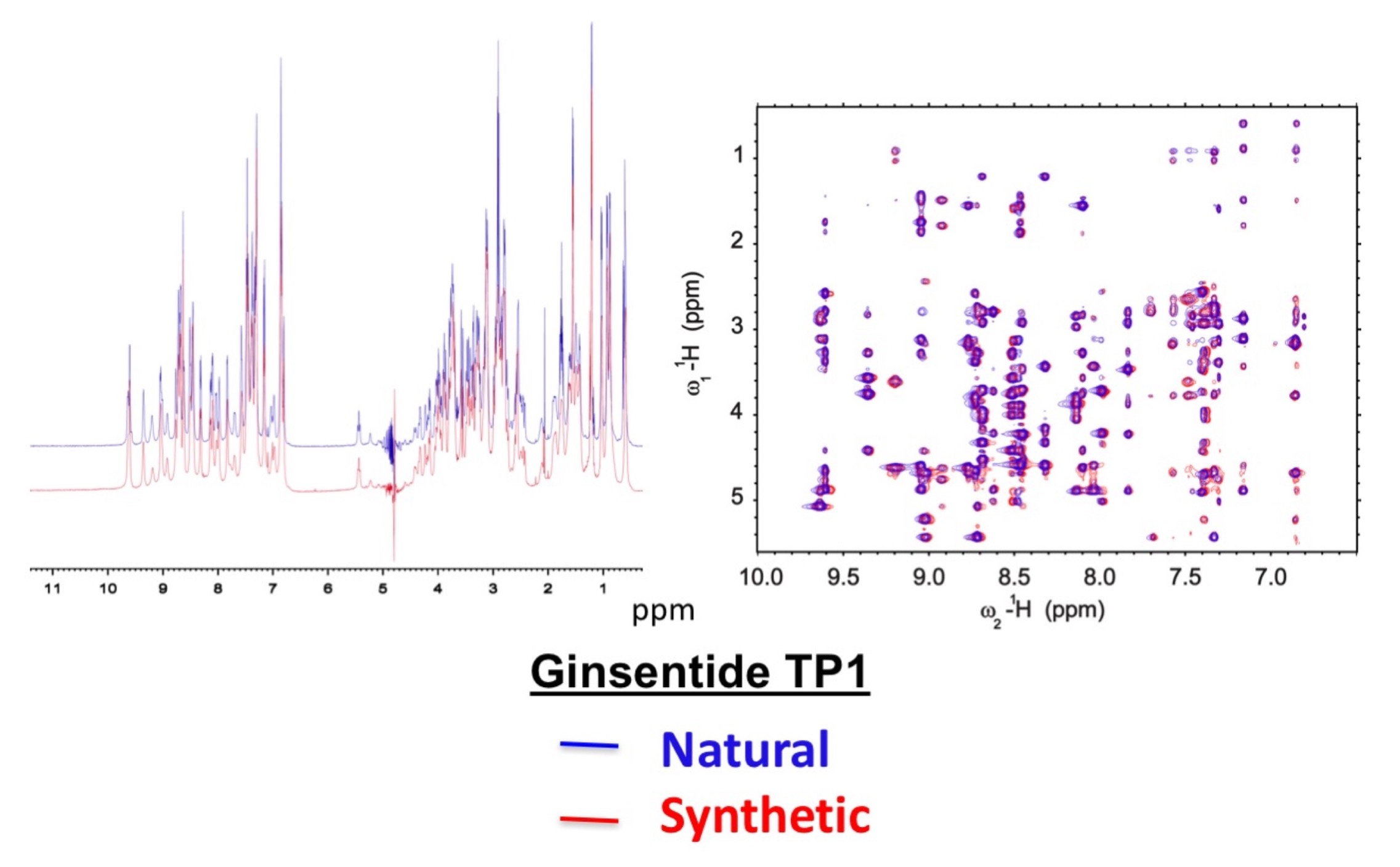
**Supplementary data S2.** Overlay of the amide-aliphatic region of natural and synthetic ginsentide TP1 by 1-dimensional and 2-dimensional nuclear Overhauser effect spectroscopy spectra.

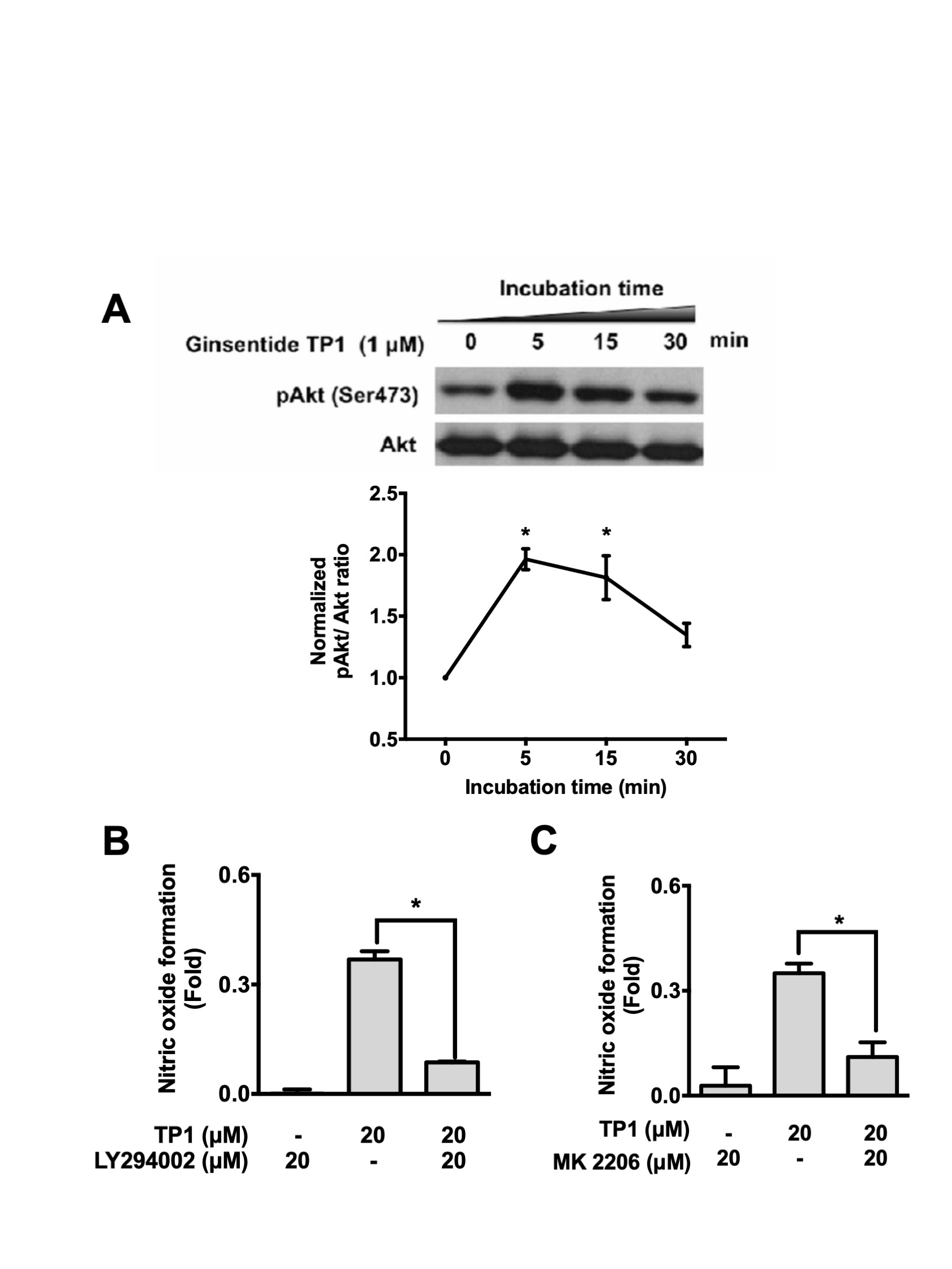

**Supplementary data S3. Ginsentide TP1-induced nitric oxide (NO) formation involves PI3K/Akt signaling.** HUVEC-CS cells were exposed to ginsentide TP1 with and without treatment of (A) MK 2206 (an Akt inhibitor) or LY294002 (a PI3K inhibitor) for 1 h. Intracellular NO production was measured with DAF-2 DA staining and expressed as normalized fluorescence intensity. (C) Representative western blot analysis of time-dependent effects of ginsentide TP1 on Akt phosphorylation in HUVEC-CS cells. All results are expressed as mean ± S.E.M. in three separate experiments. *p < 0.05 compared to the ginsentide TP1-treated group.

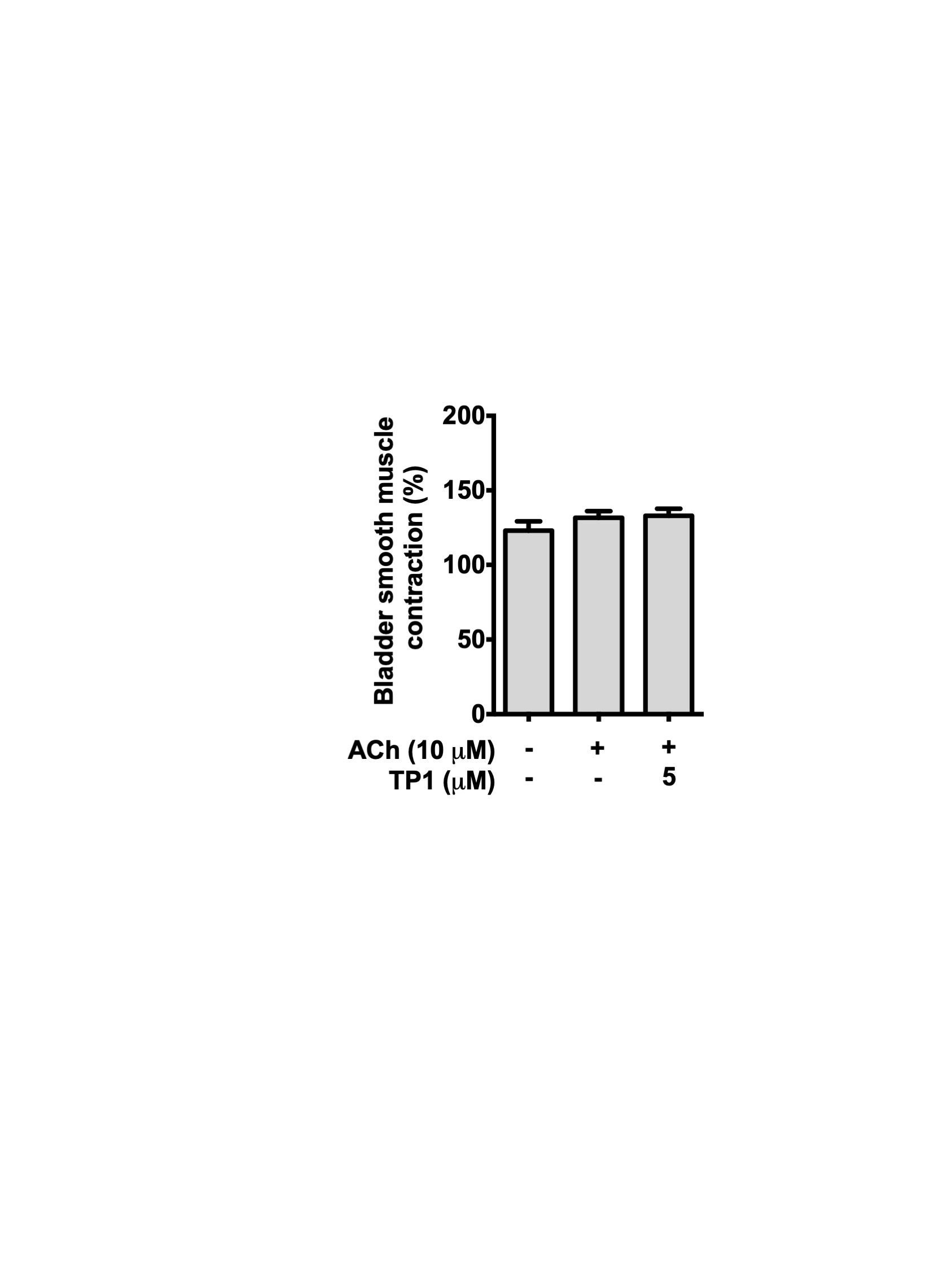

**Supplementary data S4. Ginsentide TP1 did not inhibit acetylcholine-induced bladder smooth muscle contractions.** The responses of ginsentide TP1 were characterized in acetylcholine (0.1 μM) pre-contracted bladder smooth muscle. Isometric tension was measured with a force-displacement transducer. Bladder smooth muscle contractions were measured in volts and are expressed as a percentage compared to the control. All results are expressed as the mean ± S.E.M. of two separate experiments.

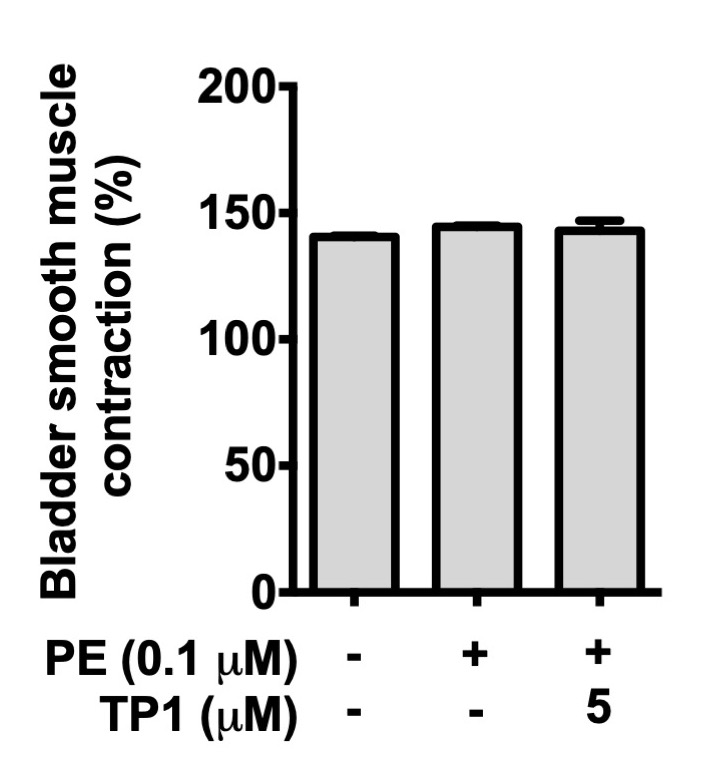

**Supplementary data S5. Phenylephrine did not induce bladder smooth muscle contractions.** Phenylephrine (0.1 μM) responses with or without ginsentide TP1 were characterized in bladder smooth muscle. Isometric tension was measured with a force-displacement transducer. Bladder smooth muscle contractions were measured in volts and are expressed as a percentage compared to controls. All results are expressed as mean ± S.E.M. of two separate experiments.

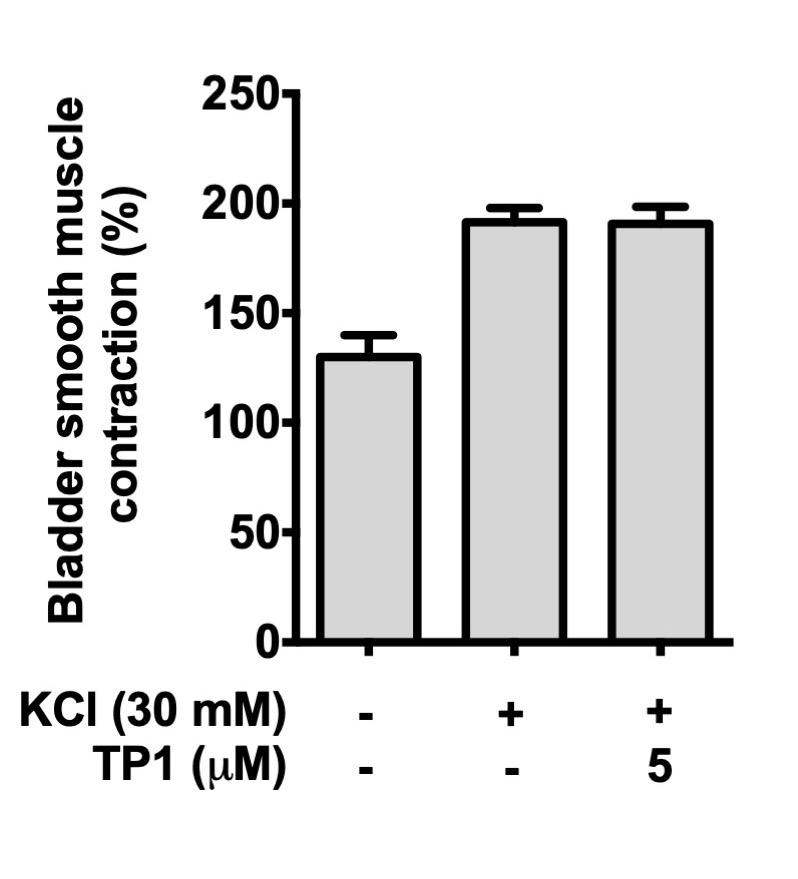

**Supplementary data S6. Ginsentide TP1 did not inhibit KCl-induced bladder smooth muscle contractions.** The responses of ginsentide TP1 were characterized in KCl (30 mM) pre-contracted bladder smooth muscle. Isometric tension was measured using a force-displacement transducer. Bladder smooth muscle contractions were measured in volts and are expressed as a percentage compared to controls. All results are expressed as mean ± S.E.M. in two separate experiments.

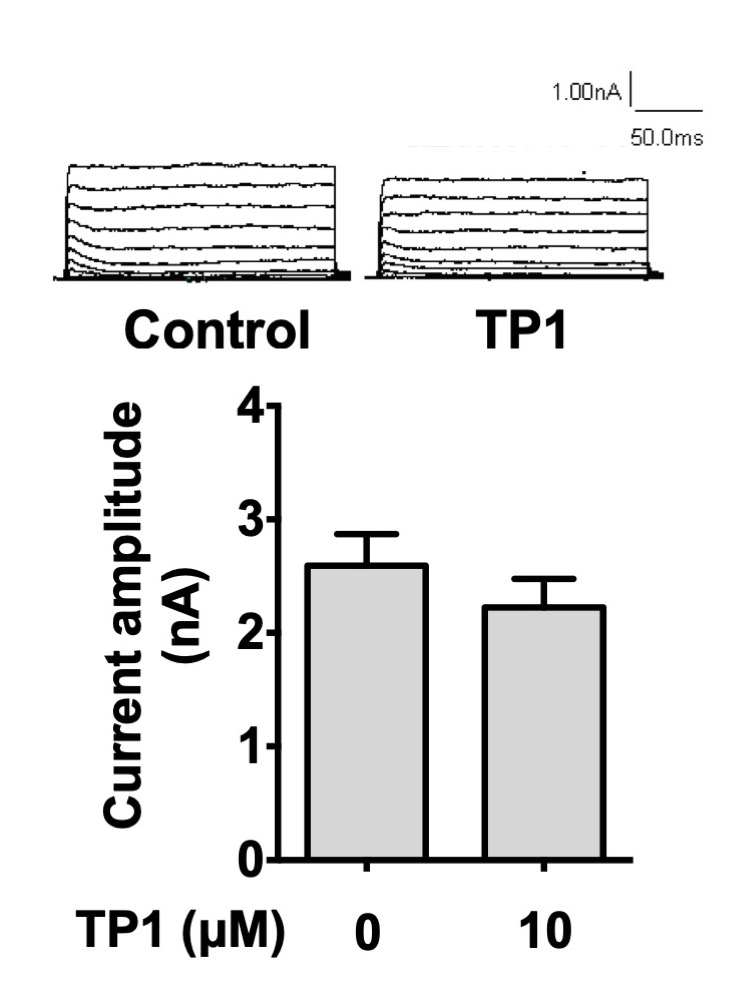

**Supplementary data S7. Ginsentide TP1 displayed no effect on the potassium current (IK) in neuronal cells.** Potassium current traces before and after the administration of 10 μM ginsentide TP1. Cells were clamped at −80 mV, followed by depolarization from −60 mV to 10 mV and further depolarization at 5 s intervals to +40 mV. This was prolonged for 200 ms to generate the total potassium current. All results are expressed as mean ± S.E.M from four separate experiments.

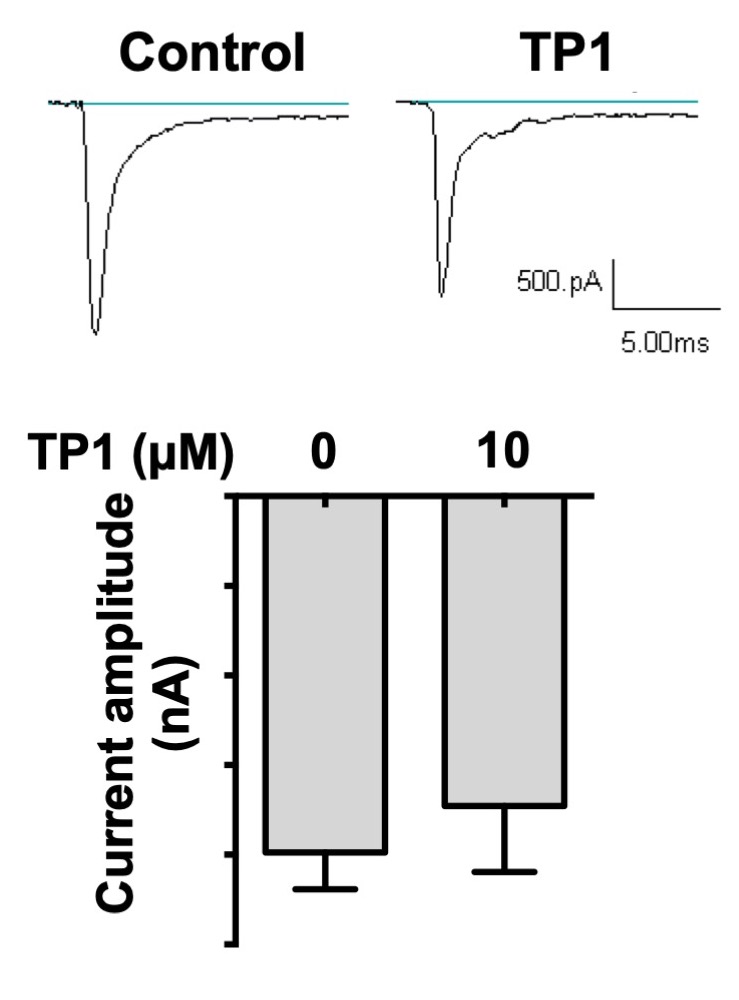

**Supplementary data S8. Ginsentide TP1 did not affect the sodium current (INA) in neuronal cells.** Sodium current traces before and after the administration of 10 μM ginsentide TP1. Cells were clamped at −100 mV followed by depolarization to −10 mV, which was prolonged for 40 ms to trigger INA. All results are expressed as mean ± S.E.M from four separate experiments.
