## Supplementary materials and methods for "Decoding the Cure-all Effects of Ginseng"

***Ex vivo* bladder smooth muscle contraction**

All protocols and procedures were approved by the Animal Care and Welfare Committee of Institute of Materia Medica, Chinese Academy of Medical Sciences and Peking Union Medical College, Beijing, China. Bladder smooth muscle contractile studies were performed as previously described with slight modifications^1^.The bladder was removed from the abdomen and two to four circular slices were cut from the bladder body after decapitation. Bladder strips weighing 15–20 mg were mounted in a double jacketed organ bath at 37 °C in Krebs solution (113 mM NaCl, 4.7 mM KCl, 1.25 mM CaCl_2_, 1.2 mM MgSO_4_, 25 mM NaHCO_3_, 1.2 mM KH_2_PO_4_, 11.5 mM glucose) and constantly bubbled with a mixture of 95% O_2_ and 5% CO_2_. The preparations were placed under initial load (1 g) and the resting tension was adjusted every 15 min. The tissues were allowed to equilibrate for 60 min and the bath fluid was changed every 15 min with fresh Krebs solution. The responses of 5 µM ginsentide TP1 were characterized in 10 µM acetylcholine, 0.1 µM phenylephrine, or 30 mM KCl pre-contracted bladder smooth muscles. . Isometric tension was measured by a force-displacement transducer and recorded using AcqKnowledge ACK100 Version 3.2 (BIOPAC Systems, USA). Bladder smooth muscle contractions were measured in volts and are expressed as a percentage compared to controls.

**Electrophysiological recordings**

**Potassium current in neuronal cells**

Effects of ginsentide on potassium currents in hippocampus neuronal cells were performed as previously described^2^. All animal experiments were approved and performed in accordance to the Animal Care and Welfare Committee of Institute of Materia Medica, Chinese Academy of Medical Sciences and Peking Union Medical College, Beijing, China. Pregnant Sprague-Dawley (SD) rats obtained from the Animal Center of Chinese Academy of Medical Sciences were killed by an overdose of isoflurane at embryonic day 18. A total of 10 fetal rats were used for hippocampus neuron dissection. The dissected uterus was placed in a sterile Petri dish and after removal of the fetuses, hippocampus tissues were obtained from their brains. The tissues were digested with 0.25% trypsin, resuspended into single cells and plated on poly-D-lysine–coated coverslips. The cells were maintained in culture medium consisting of neurobasal medium with B-27 supplement (Invitrogen, USA), penicillin, streptomycin and 2 mM L-glutamine. Day 10-14 cultured neuronal cells were used for electrophysiological recording.

Traditional whole-cell voltage-clamp recordings were performed at room temperature (22–25 °C) to record the potassium currents. Recording pipettes were pulled with borosilicate glass (World Precision Instruments, Inc., Sarasota, FL, USA) to 2–5 MΩ when filled with a pipette solution containing 145 mM KCl, 1 mM MgCl_2_, 5 mM EGTA, 10 mM HEPES and 5 mM MgATP at pH 7.2 and placed in a bath solution containing 140 mM NaCl, 5 mM KCl, 2 mM CaCl_2_, 1.5 mM MgCl_2_, 10 mM glucose and 10 mM HEPES at pH 7.4 adjusted with NaOH. Isolated cells were voltage clamped in whole-cell mode with an AxoPatch 200B amplifier (Molecular Devices Corporation, Sunnyvale, CA, USA), and currents were recorded at 10 kHz. Cells were continuously perfused with bath solution through a gravity-driven perfusion system (ALA Scientific, Farmingdale, NY, USA). Potassium current traces before and after the administration of 10 μM ginsentide TP1 was measured. Cells were clamped at −80 mV, followed by depolarization from −60 mV to 10 mV and further depolarization at 5 s intervals to +40 mV. This was prolonged for 200 ms to generate the total potassium current. Electrophysical data were processed in Clampfit 9.2 (Molecular Devices,Sunnyvale, CA, USA) and then analysed in Excel and Origin 6.0 (OriginLab Corporation, Northampton, MA, USA).

**Sodium current in neuronal cells**

Effect of ginsentide TP1 on sodium currents in dorsal root ganglion (DRG) neurons was examined as previously described^3^. DRG neurons from adult male SD rats obtained from the Animal Center of Chinese Academy of Medical Sciences were used to record neuronal sodium currents. All experiments were approved and performed in accordance to the Animal Care and Welfare Committee of Institute of Materia Medica, Chinese Academy of Medical Sciences and Peking Union Medical College, Beijing, China. Adult male SD rats were killed with an overdose of isoflurane. DRGs were collected in cold DH10 [90% DMEM/F-12 (Gibco, USA), 10% FBS (Gibco, USA), 1% penicillin-streptomycin] and then treated with enzyme solution [3.5 mg/ml dispase, 1.6 mg/ml collagenase type I, and 1 U/ml DNase in HBSS (Hanks’ balanced salt solution) (Gibco, USA) without Ca^2+^ and Mg^2+^] at 37 °C. After centrifugation, dissociated cells were resuspended in DH10 and plated at a density of 1.5 X 10^5^- 4 X 10^5^ cells on glass coverslips coated with poly-L-lysine (0.5 mg/mL; Sigma-Aldrich, USA) and laminin (10 mg/mL; Invitrogen, USA). The cells were cultured in Neurobasal-A Medium (Gibco, USA); supplemented with B27 (Gibco, USA), NGF (Nerve growth factor), and GlutaMAX. After 2 h dissociation, the cells were used for electrophysiological recording using manual patch-clamp.

To record sodium channel currents from DRG neurons, whole-cell voltage-clamp recording was performed at room temperature. Recording pipettes were pulled with borosilicate glass to ∼1–2 MΩ for DRG sodium channels. To reduce voltage errors, low-sodium external solution (35 mM NaCl, 105 mM choline-Cl, 1 mM CaCl_2_, 1 mM MgCl_2_, 20 mM tetraethylammonium, 0.1 cadmium-Cl, 10 mM glucose, and 10 mM HEPES at pH 7.4 adjusted with NaOH) was used to record total sodium currents. Internal solution for sodium channels contained 140 mM CsF, 10 mM NaCl, 1 mM EGTA, and 10 mM HEPES at pH 7.3 adjusted with CsOH. Cells were voltage-clamped in whole-cell mode with an EPC-10 amplifier (HEKA, Lambrecht/Pfalz, Germany), and currents were sampled at 10 kHz. Cells were continuously perfused with external solution through a gravity driven perfusion system (ALA Scientific, Farmingdale, NY, USA). Sodium current traces before and after the administration of 10 μM ginsentide TP1 were measured. Cells were clamped at −100 mV followed by depolarization to −10 mV, which was prolonged for 40 ms to trigger sodium current. Electrophysiological data were processed in FitMaster (HEKA, Lambrecht/Pfalz, Germany) and analyzed in Excel (Microsoft, Redmond, WA) and Origin 6.0 (Origin Lab, Northampton, MA).
